# Hyaluronic Acid and Emergent Tissue Mechanics Orchestrate Digit Tip Regeneration

**DOI:** 10.1101/2024.12.04.626830

**Authors:** Byron W.H. Mui, Joseph Y. Wong, Toni Bray, Lauren Connolly, Jia Hua Wang, Alexander Winkel, Pamela G. Robey, Kristian Franze, Kevin J. Chalut, Mekayla A. Storer

## Abstract

Tissue regeneration is a dynamic process requiring coordinated cell fate decisions to restore structure and function. For instance, the tips of human and rodent digits fully regrow after amputation, although more proximal injuries fail to regenerate. While biochemical cues have been widely studied in wound healing, the role of the physical microenvironment remains less understood. Here, we discovered that tissue mechanics and extracellular matrix (ECM) composition differ markedly between non-regenerating and regenerating wounds. These differences are due to specific fibroblast sub-types which predominate in the non-regenerating wound and form fibrotic collagen networks. Conversely, bone-forming progenitors in regenerating digits synthesize abundant hyaluronic acid, which mediates collagen fibrillogenesis. By modeling the tissue mechanics of non-regenerative and regenerative wounds using hydrogels, we showed that substrate stiffness regulates the synthesis of fibrotic and regenerative ECM. We further revealed that mimicking the regenerative wound mechanics upregulated bone morphogenic protein signaling and osteogenic differentiation. We ultimately demonstrated that the link protein HAPLN1 promotes hyaluronic acid deposition *in vivo*, reduces scar formation, and induces bone repair after non-regenerative amputations. These findings emphasize the interplay among the ECM, tissue mechanics, and cell behavior in digit regeneration and point towards ECM modulation as a strategy to improve wound healing.

**Highlights:**

- Non-regenerating and regenerating digit tips diverge in their cellular and ECM composition
- Regeneration requires hyaluronic acid matrix, which mediates collagen assembly and tissue mechanics
- Soft substrates enhance BMP signaling and propagate the synthesis of regenerative ECM
- HAPLN1 overexpression initiates rescue of non-regenerative digit tip amputations

## Introduction

Adult injuries often heal through scar formation, characterized by the deposition of connective tissue known as fibrosis. Although normal wound healing relies upon the synthesis of extracellular matrix (ECM) as provisional scaffolding, excessive accumulation of ECM may disrupt tissue architecture and lead to substantial disease burden^1,2^. Developing effective treatments to halt or reverse fibrosis in favor of regeneration hinges on advancing our understanding of wound healing^3,4^. To that end, amputation of mouse digits presents a unique opportunity to examine mammalian regeneration and fibrosis in parallel^5,6^. Severing the tip of the digit results in complete, multi-tissue restoration^5,6^. Central to digit regeneration is the blastema^7^, an intermediate structure harboring heterogeneous progenitors^8,9^ that restore lost tissues^10,11^. More proximal amputations, such as those that sever the second phalanx, fail to regenerate^5^. Instead, they exhibit severely stunted or no regrowth, and dense fibrous connective tissue accumulates at the injury site^5^. While previous studies have characterized cellular and molecular machineries behind digit tip wound healing^8,9,12–17^, few have examined the role of the ECM^18^. Thus, in the adult digit amputation model, we investigated how the ECM may drive regeneration instead of fibrosis.

The ECM is a complex and dynamic network of fibrous proteins and proteoglycans that provides structural support and signaling to cells within tissue^19^. Evidence mounts that the ECM provides essential chemical and mechanical information within various biological contexts, including tissue morphogenesis^20–22^, homeostasis^23,24^, aging^25,26^, and cancer^27–30^. Studies have also shown how the ECM impacts regeneration^31,32^ or fibrosis^33^. For example, early studies on scarless fetal cutaneous injuries implicated a protective role for the ECM component hyaluronic acid (HA), a linear, non-sulfated polymer composed of repeating disaccharide units of *N*-acetyl-glucosamine and glucuronic acid^34^. Indeed, fetal wounds remarkably heal with minimal collagen deposition while sustaining high levels of HA, whereas adult wounds show extensive infiltration of collagen concurrent with rapid degradation of HA^35–37^. A further study showed that HA depletion in fetal wounds induces fibrosis^38^. More recent studies have similarly proposed favorable roles for HA in regeneration in other animal species^39–41^ or mammalian organ systems^42,43^. However, in adult mammals, the influence of HA on regeneration and its relationship to fibrosis remains poorly understood.

Mechanical cues from the niche are potent regulators of cellular behavior^44–48^. In fibrotic wound repair, the ECM typically stiffens^49,50^, which mechanically activates fibroblasts^51–53^, the main cellular regulators of ECM. Importantly, other mechanical properties of ECM and how they drive fibroblast response have been largely overlooked, such as viscosity^54^. Nevertheless, in response to ECM changes, fibroblasts further remodel their extracellular environment^55,56^. In other words, changes in the ECM composition^57^ or organization^58–60^ alter the overall tissue mechanics, which feeds back to mediate fibroblast activity. Processes such as apoptosis^61,62^ or inactivation^63^ of fibroblasts return wound repair programs to homeostasis. However, through largely unknown mechanisms, fibrosis progression bypasses these pathways, leading to cumulative disruption of the tissue architecture. Given that fibroblasts are heterogeneous^64–67^, some evidence suggests an intrinsic propensity for fibrosis by specific sub-types of fibroblasts^68,69^, which may be mostly absent from regenerative tissues. Simultaneously, it is also possible that fibrosing and regenerating ECM niches are distinct and enable opposing regulatory mechanisms. With the goal of addressing these unknowns, we investigated how cell-derived ECM relate to emergent tissue mechanics and influence digit wound healing.

In this study, we interrogated the divergence between non-regenerating and regenerating digits following amputation. We show that tissue mechanics, alongside cell and ECM composition, differentiate digit repair programs and highlight ways in which the ECM plays an active role in wound healing. Lastly, we provide evidence for ECM modulation as a viable strategy to promote regeneration.

## Results

### Collagen underlies tissue stiffness in amputated digits

Fibrous tissue accumulates after non-regenerative amputations^5^, but little is known about the blastema’s ECM. Therefore, we used second harmonic generation microscopy to investigate collagen networks^70^ after digit amputations. Compared to the blastema, non-regenerative wounds contained more collagen (Figures 1A, 1B, S1A, and S1B). Additionally, collagen fibers were wider, longer, and more aligned (S1C-1E). The elevated collagen content and increased structural maturity observed in non-regenerative wounds confirmed the presence of fibrotic ECM, an absent feature in the blastema. Given the importance of collagen in tissue mechanics^71^, we used atomic force microscopy (AFM) to investigate the physical properties of wound digits. We detected mechanical differences between non-regenerative wounds and the blastema, including three-fold greater stiffness in the former (Figures 1C, 1D, S1F, and S1G). Collectively, these results suggest that the non-regenerative and regenerative wounds have very different underpinning ECM microenvironments.

**Figure 1.**
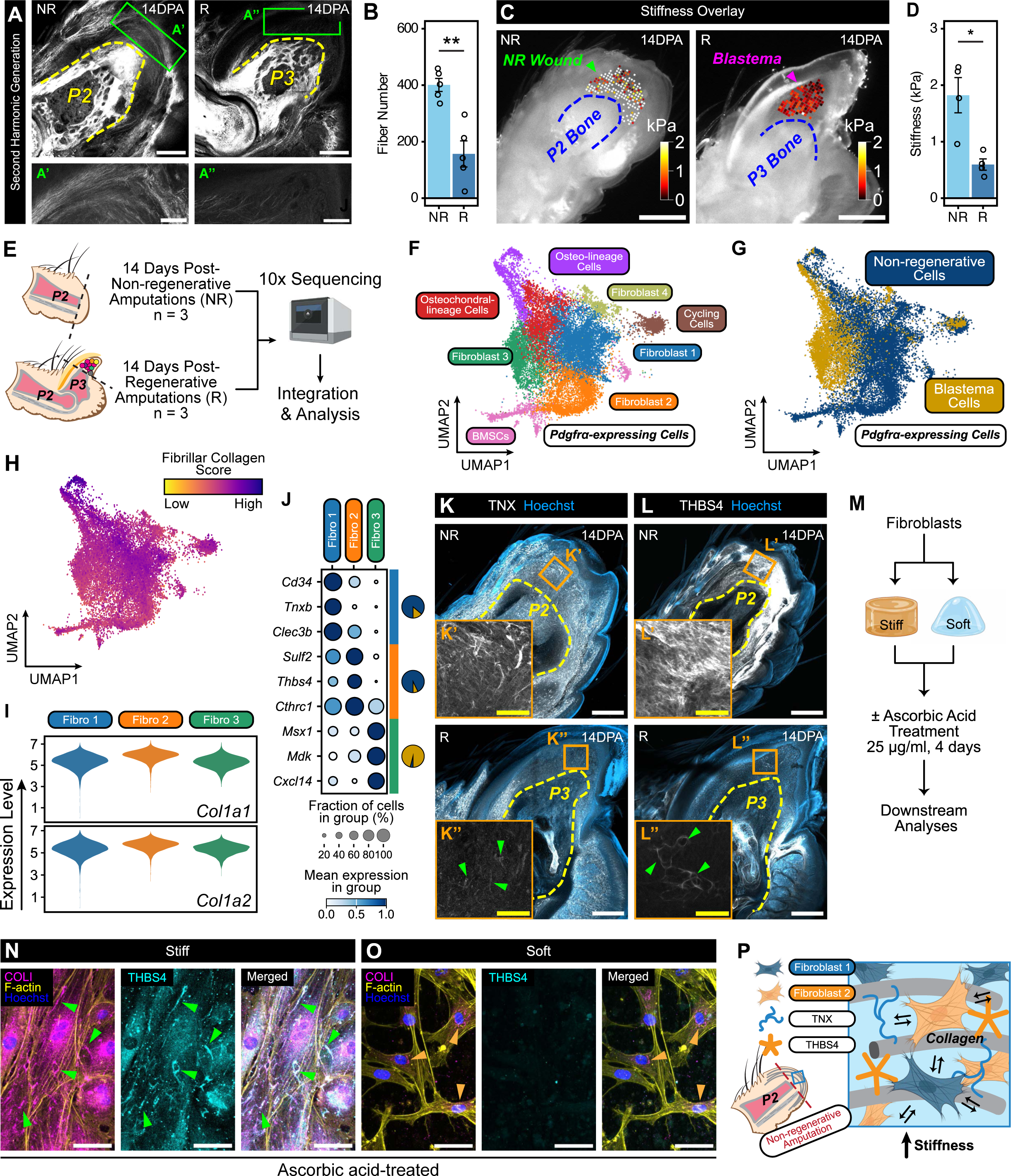
Collagen remodeling propagates tissue stiffness in non-regenerating digits. (A and B) Second harmonic generation microscopy images of non-regenerating (NR) and regenerating (R) digits 14 days post-amputation (DPA), highlighting collagen I fibers. The dashed yellow line indicates the border of the second (P2) or third (P3) phalanx bone. Scale bars = 250 µm. The bottom panels show magnified views of the green boxed regions of fibrosing (A’) and blastema (A”) tissues. Scale bars = 100 µm. Fiber number was quantified in (B). n = 5 mice per condition. (C and D) Atomic force microscopy stiffness maps of the 14DPA NR wound (green arrowhead) and blastema (magenta arrowhead). The dashed blue line indicates the border of the P2 or P3 bone. Scale bars = 500 µm. Average stiffnesses were quantified in (D), which were determined by applying the Hertz model. n = 4 mice per condition. (I) (E) Schematic of the single-cell RNA sequencing experiment of 14DPA NR and R digit tissues. Two 14DPA non-regenerative datasets were original datasets generated from this paper. n = 2 replicates of 4 mice per replicate. The remaining datasets were acquired from the public domain. n = 1, 14DPA non-regenerative dataset (GEO: GSE135985); n = 3, 14DPA blastema datasets (GEO: GSE135985 and GSE143888). (F-H) Uniform Manifold Approximation and Projection (UMAP) of 20,536 *Pdgfrα*-expressing cells from 14DPA NR and blastema datasets. In (F), cells are color-coded according to clusters, which include four fibroblast (Fibroblast 1, 2, 3, and 4), osteochondral-lineage, osteo-lineage, cycling, and bone marrow stromal cell (BMSC) clusters. In (G), cells are color-coded by NR and blastema datasets. In (H), each cell was assigned a score based upon its expression of fibrillar collagen genes, where cells with a low score appear as yellow, and a high score dark purple. (I) Violin plots of *Col1a1* and *Col1a2* expression among Fibroblast 1 (Fibro 1), Fibroblast 2 (Fibro 2), and Fibroblast 3 (Fibro 3) cells. (J) Dot plot showing top differentially expressed genes for each fibroblast sub-type. The size of the circle represents the fraction of cells expressing the gene, and average gene expression is shown according to the key, in which dark blue and white represent high and low expression, respectively. Pie charts represent the relative proportion of NR (blue) and blastema (gold) cells within a specific fibroblast sub-type. (K and L) Immunofluorescence images of 14DPA NR and R digits stained for TNX (white) in (K) and THBS4 (white) in (L). The yellow dashed line indicates the border of the P2 or P3 bone. All images were counterstained with Hoechst (blue). Scale bars = 250 µm. Insets show the orange boxed regions of fibrosing (K’ and L’) and blastema (K” and L”) tissues at higher magnification. Green arrowheads indicate dim TNX and THBS4 expression in the blastema. Scale bars = 50 µm. (M) Schematic of the experimental design to test the effects of substrate stiffness on fibrotic ECM synthesis. Ascorbic acid was supplemented to induce collagen fibrillogenesis. (N and O) Immunofluorescence images of fibroblasts cultured on stiff (50 kPa; N) or soft (0.7 kPa; O) hydrogels, treated with ascorbic acid, and stained for COLI (magenta), F-actin (yellow), THBS4 (cyan), and Hoechst (blue). In (N), green arrowheads point at collagen fibers co-localizing with THBS4. In (O), orange arrowheads point at intracellular collagen. Scale bars = 50 µm. (P) Working model of the cellular and ECM composition of the wounded P2 digit after non-regenerative amputations. Bidirectional arrows indicate cell-ECM feedback. Data are shown as mean ± SEM. Statistical significance was determined by two-tailed unpaired student’s *t*-test. *p < 0.05, **p<0.01, ***p<0.001, ns = not significant.

### Distinct fibroblast sub-types producing collagen remodeling glycoproteins dominate non-regenerative wound healing

Given the clear differences in collagen structures and tissue stiffness between non-regenerating and regenerating digits, we next investigated what factors may be underlying the fibrotic response. To do this, we conducted single-cell analysis of non-regenerative wounds and integrated these datasets with blastema datasets as controls (Figure 1E). Unsupervised clustering yielded seven distinct cell populations (Figures S1H and S1I). Given that a hallmark of fibrosis is excessive ECM accumulation, our first objective was to identify the primary cell type responsible for establishing the ECM. Each cell was assigned an ECM score based upon its expression of core matrisome^72^ genes (Figure S1J). The *Pdgfrα*-expressing cluster showed the highest ECM activity, outperforming all other clusters (Figure S1J). Gene set enrichment analysis revealed that this cluster upregulated ECM-related pathways, including ‘Collagen Fibril Organization’ and ‘Extracellular Matrix Organization’ (Figure S1K). We deduced that fibroblasts—the main cell type involved in establishing the extracellular milieu^73^—likely comprised the *Pdgfrα*-expressing cluster. We focused on this cell population to explore how it might promote fibrosis^74^ and counteract the regenerative process.

Fibroblast heterogeneity exists within and across organs^66,75^. As such, we hypothesized that wound ECMs diverge due to differences in cellular composition. Sub-setting *Pdgfrα-*expressing cells revealed seven distinct transcriptional signatures: four fibroblast clusters denoted Fibroblast 1, Fibroblast 2, Fibroblast 3, and Fibroblast 4; osteochondral-lineage; osteo-lineage (OL); cycling; and bone marrow stromal cell clusters (Figure 1F). Since we observed only a modest overlap between *Pdgfrα*-expressing cells from the non-regenerative wound and blastema (Figure 1G), we queried whether specific fibroblast sub-types are preferentially enriched in one wound healing condition over the other. To investigate this, we analyzed the proportion of each cell type within non-regenerative and regenerative conditions. Our analysis showed that Fibroblast 1 and 2 cells were the predominant sub-types in the former, while Fibroblast 3 cells were the primary sub-type in the latter (Figure S1L). Furthermore, a correlation analysis of average gene expressions showed that a fibroblast sub-type from the non-regenerative tissue was not transcriptionally distinct from the same sub-type in the blastema (Figure S1M). Taken together, the predominance of fibroblast sub-types may be a stronger determinant of the wound response than changes in the activity of the same cell type across conditions.

Since Fibroblast 1 and 2 cells were most prevalent in non-regeneration, we hypothesized that their activity accounts for the fibrogenic response. All *Pdgfrα*-expressing clusters expressed fibrillar collagens to a similar degree (Figures 1H and 1I), suggesting that collagen fibrillogenesis is mediated post-translationally. We identified elevated synthesis of tenascin X (TNX)—which promotes collagen packing^76,77^—and CD34^+^ Fibroblast 1 cells in the non-regenerative wound, both mostly absent in the blastema (Figures 1J, 1K, and S1N). Similarly, in the non-regenerative wound, Fibroblast 2 cells produced high levels of thrombospondin-4 (THBS4)—a factor that maintains the ECM in tendon and myotendinous junctions^78^—while the blastema exhibited low expression (Figures 1J and 1L). Of note, Fibroblast 3 cells, the major blastema cell type, did not differentially express major collagen remodeling genes (Figure 1J). Instead, they expressed markers associated with limb regeneration, such as *Msx1*^79^ and *Mdk*^80,81^ (Figure 1J). Taken together, Fibroblast 1 and 2 cells—which upregulated TNX and THBS4, respectively—were specific to non-regeneration, suggesting that they drive the fibrotic response through collagen remodeling.

### Elevated stiffness propagates the synthesis of pro-fibrotic ECM

Given that fibroblasts are highly mechanosensitive and are central players in ECM remodeling^74^, we hypothesized that non-regenerative ECM is reinforced through feedback mechanisms involving tissue mechanics. To test this hypothesis, we fabricated stiff and soft StemBond hydrogels^82^ to model the mechanical microenvironment of the non-regenerative wound and the blastema, respectively (Figure 1M). Using ascorbic acid to induce collagen synthesis^83^, we asked how stiffness influences collagen I fibrillogenesis and THBS4 synthesis, both of which typified non-regenerative healing (Figure 1M). Fibroblasts cultured in stiff conditions exhibited cell spreading and developed prominent actin cytoskeleton (Figures 1N and S1O). After ascorbic acid treatment, fibroblasts produced fibrillar collagen networks, which largely co-localized with THBS4 (Figure 1N). Conversely, in soft conditions that mimic the blastema, collagen I-THBS4 network formation remained low, irrespective of ascorbic acid addition (Figures 1O and S1P). These results suggested a self-sustaining feedback loop in which substrate stiffening enhances the synthesis of fibrosis-associated ECM that further stiffens the environment (Figure 1P).

### Osteo-lineage cells are the primary ECM synthesizers in regeneration

Given that fibrosis is the antithesis of regeneration, we hypothesized that the blastema is enriched with cell sub-types, distinct from Fibroblast 1 and 2 cells, that are responsible for synthesizing regeneration-specific ECM. Our cell sub-type correlation analysis demonstrated that Fibroblast 1 and 2 cells were most disparate from osteo-lineage cells (OLs) (Figure S1M), which was roughly three times more prevalent in the blastema (Figure S1L). Supporting this distinction, comparisons of Fibroblast 1 and 2 clusters against all other *Pdgfrα*-expressing cells showed that both sub-types of fibroblasts exhibited prominent downregulation of osteoblast-related genes *Alpl*, *Spp1*, and *Ibsp* (Figures S2A and S2B). Lastly, ECM scoring indicated that OLs had the highest expression of collagens and proteoglycans (Figure S2C). Altogether, we posited that while Fibroblast 1 and 2 cells are drivers of the non-regenerative ECM, the blastema’s main ECM-modulating counterpart are the osteo-lineage cells (OLs).

### Hyaluronic acid, aggrecan, and link protein accumulate within the blastema

We then investigated OL-synthesized ECM components. We performed pathway and differential gene expression analyses, comparing OLs against Fibroblast 1 and 2 cells separately (Figures 2A and S2D). The emergence of terms such as “Glycosaminoglycan Binding”, “Proteoglycan Binding,” and “Hyaluronic Acid Binding” (Figures 2B and S2E) suggested an OL microenvironment in which non-collagenous matrisome components, particularly hyaluronic acid (HA), appear prominently in the blastema. OLs expressed *Runx2*, signifying their fate as committed osteogenic cells, as well as *Id1*, an early target gene of BMP signaling^84^ (Figures 2C and S2F). Although P3 bone regenerates without the formation of a cartilage callus, OLs highly expressed the transcription factors *Sox6* and *Sox9*^85^, along with cartilage-specific target genes *Acan* and *Col2a1* (Figures 2C, S2F, and S2G). In conjunction, OLs upregulated *Hapln1* (Figures 2C, S2F, and S2G). HAPLN1 links aggrecan to HA^86^, conferring structural integrity to the overall HA-HAPLN1-ACAN complex^87,88^. From the transcriptomic data, we deduced that the blastema’s ECM starkly contrasts non-regenerative ECM through higher representation of non-collagenous, OL-derived ECM elements comprising HA and its binding partners HAPLN1 and ACAN.

**Figure 2.**
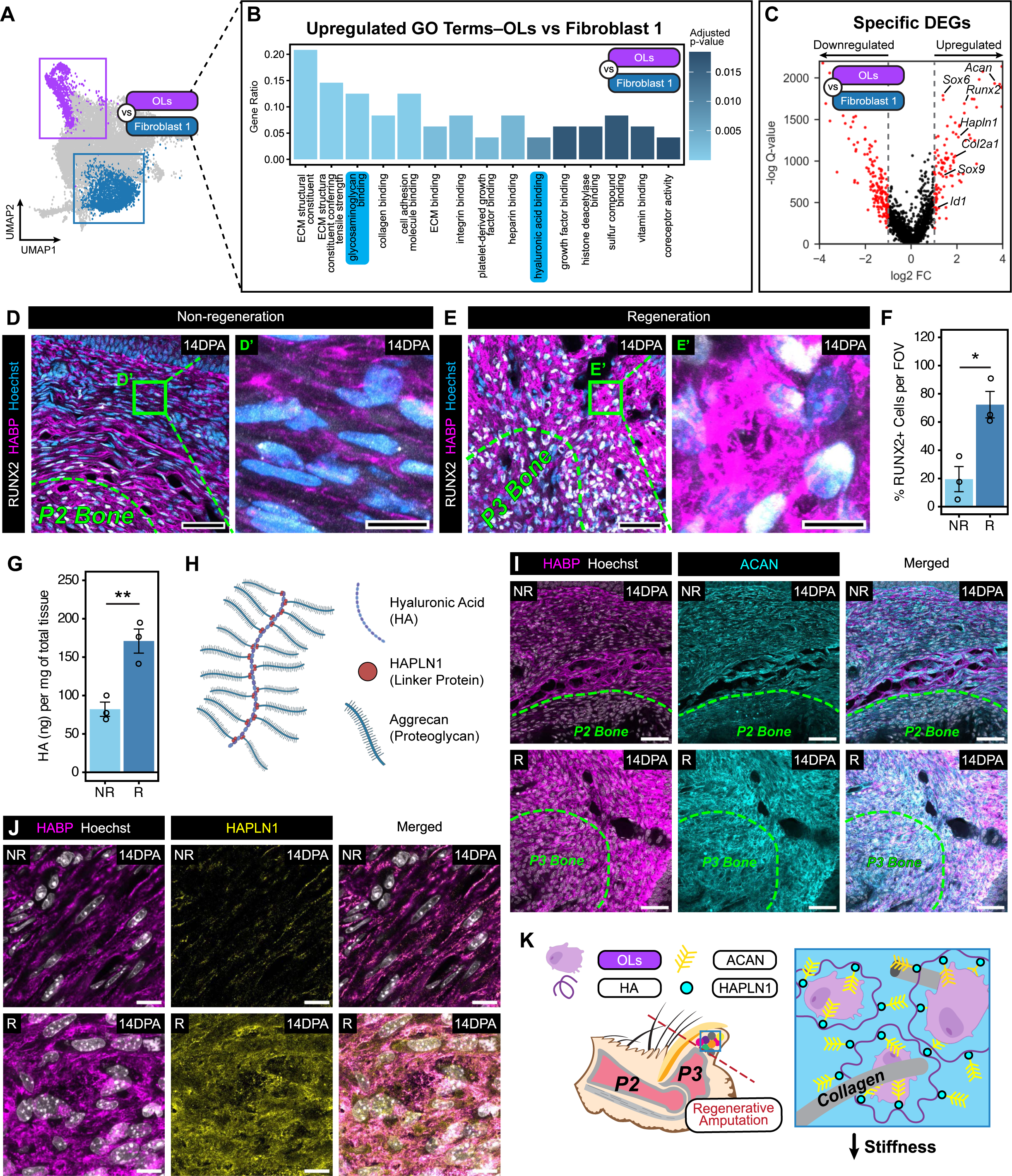
Hyaluronic acid is central to the blastema’s regenerative ECM. (A) UMAP diagram showing the comparison of the osteo-lineage cluster (OLs, purple) against the Fibroblast 1 cluster (dark blue). Boxed regions are highlighting the clusters of interest. (B) Gene ontology terms upregulated in osteo-lineage cells (OLs) compared to Fibroblast 1 cells, determined by gene set enrichment analysis using clusterProfiler. The Gene Ratio indicates the proportion of input genes associated with a gene ontology term, and the color indicates the adjusted p-value of the enrichment score. (C) Volcano plot of differentially upregulated genes in osteo-lineage cells (OLs) compared to Fibroblast 1 cells, determined by the two-part generalized linear model MAST. Genes related to hyaluronic acid and chondrogenesis are annotated. Thresholds were set at -1 > log2FC > 1 and Q-value < 0.05. (D-F) Immunofluorescence images of 14 days post-amputation (DPA) non-regenerating (NR; D) and regenerating (R; E) digits stained for RUNX2 (white), HABP (magenta), and Hoechst (blue). The green dashed line indicates the border of the second (P2) or third (P3) phalanx bone. Scale bars = 50 µm. The right-hand panels show magnified views of the green boxed regions of fibrosing (D’) and blastema (E’) tissues. Scale bars = 10 µm. The percentage of RUNX2^+^ cells was quantified in (F). n = 3 mice per condition. (G) Quantification of total HA per mg of 14DPA NR and R tissues by enzyme-linked immunosorbent assay. n = 3 replicates per condition of 4 mice pooled per replicate. (H) Diagram of HA-HAPLN1-ACAN complex. (I and J) Immunofluorescence images of 14DPA NR (top panel) and R (bottom panel) digits stained for HABP (I and J; magenta), ACAN (I; cyan), and HAPLN1 (J; yellow). Sections were counterstained with Hoechst (white). In (I), the green dashed line indicates the border of the P2 or P3 bone. Scale bars in (I) = 50 µm, and in (J) = 10 µm. (K) Working model of the cellular and ECM composition of the blastema after regenerative amputations. OLs = osteo-lineage cells, HA = hyaluronic acid, ACAN = aggrecan, and HAPLN1 = hyaluronic acid and proteoglycan link protein 1. Data are shown as mean ± SEM. Statistical significance was determined by two-tailed unpaired student’s *t*-test. *p < 0.05, **p<0.01, ***p<0.001, ns = not significant.

We proceeded to investigate, *in situ*, whether OLs are more prevalent in regeneration, and if this corresponded to a greater accumulation of hyaluronic acid (HA). RUNX2^+^ OLs were globally diminished in the non-regenerative wound (Figures 2D and 2F), coincident with sparse HA (Figures 2D, S2H-S2J). In comparison, the blastema was densely populated by RUNX2^+^ OLs encased in abundant HA matrix, reminiscent of pericellular coats or glycocalyx^89^ (Figures 2E, 2F, S2H-S2J). These data suggested that OLs aggregate HA in their local environment. By performing enzyme-linked immunosorbent assay of HA extracted from non-regenerative and blastema tissues, we confirmed that total HA quantity was two-fold greater in the blastema (Figure 2G). We then assayed the distribution of HAPLN1 and ACAN, which decorates HA in a brush-like configuration^90^ (Figure 2H). ACAN and HA were diffusely expressed at low levels in the non-regenerative wound (Figure 2I). In contrast, the blastema demonstrated a strong presence of ACAN that co-localized with HA (Figure 2I). Moreover, unlike in the non-regenerative wound, an abundance of HA and HAPLN1 overlapped in the blastema (Figure 2J). Altogether, the blastema’s OLs synthesize copious matrices composed of HA, HAPLN1, and ACAN in the absence of major collagen networks, forming a distinct ECM niche in regeneration (Figure 2K).

### Depletion of hyaluronic acid impairs bone regrowth and blastema formation

Next, we asked whether hyaluronic acid (HA) actively contributes towards wound healing. To answer this question, we enzymatically degraded HA by serial injections of hyaluronidase into the digit tip after regenerative amputations (Figure 3A). At 14DPA, digits were decreased in length, and, though we did not find the difference to be statistically significant, the area was often smaller (Figures 3B, 3C, and S3A). RUNX2^+^ osteo-lineage cells (OLs) and pericellular HA were also diminished after hyaluronidase treatment, corroborating the relationship between OLs and HA (Figures 3D-3F).

**Figure 3.**
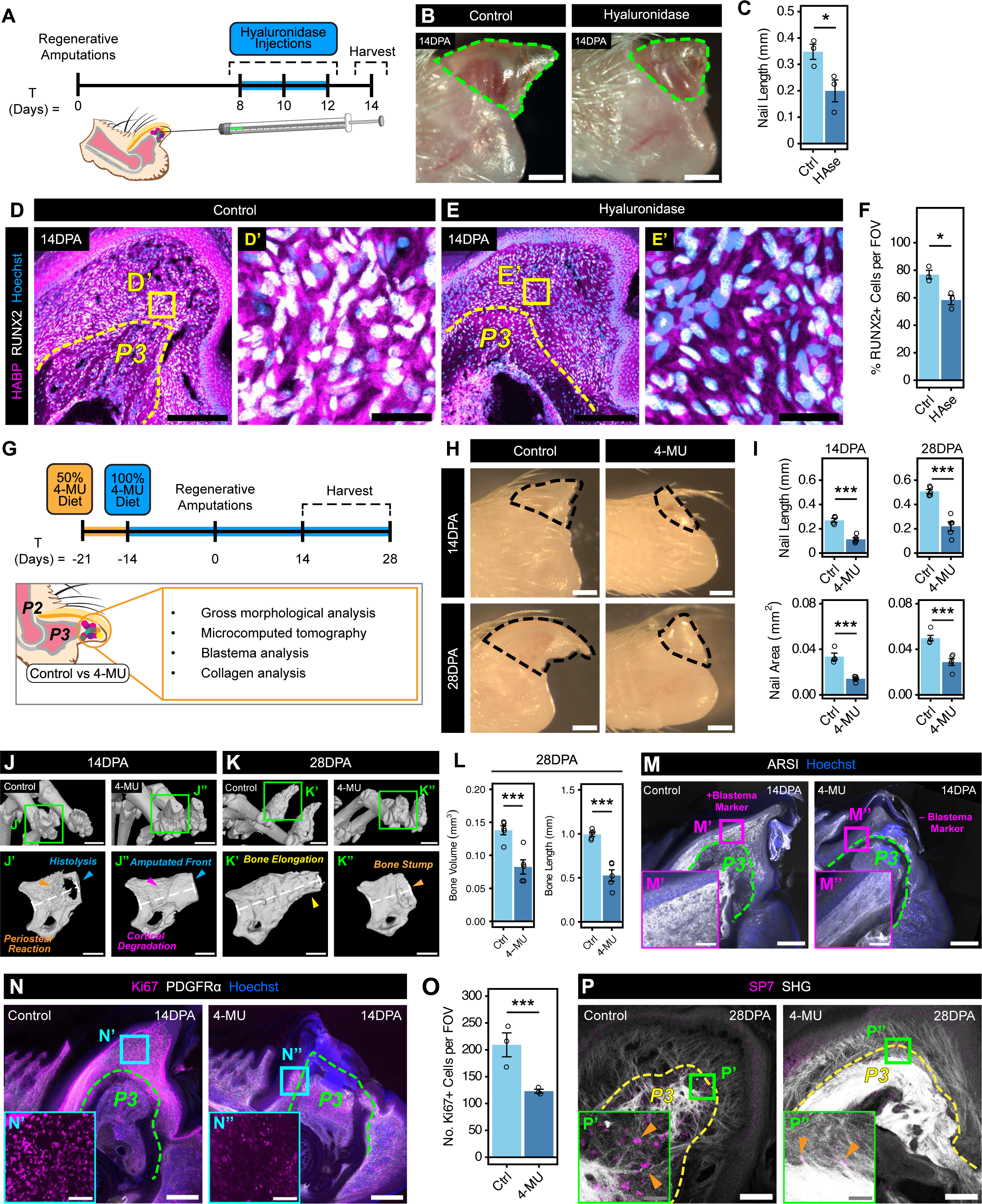
Digit tip regeneration requires hyaluronic acid. (A) Schematic of the experimental design using hyaluronidase to degrade hyaluronic acid (HA) after regenerative amputations. (B and C) Gross images of control (left) or hyaluronidase (right) digits at 14 days post-amputation (DPA). The green dashed line indicates the border of the third phalanx (P3) nails, which were assessed for length in (C). Scale bars = 100 µm. n = 3 mice per condition. (D-F) Immunofluorescence images of 14DPA control (D) or hyaluronidase (E) digits stained for HABP (magenta), RUNX2 (white), and Hoechst (blue). The yellow dashed line indicates the border of the P3 bone. Scale bars = 200 µm. The right-hand panels (D’ and E’) show the yellow boxed regions at higher magnification. Scale bars = 30 µm. The percentage of RUNX2^+^ cells per FOV was quantified in both conditions in (F). n = 3 mice per condition. (G) Schematic of the experimental design testing continuous knockdown of hyaluronic acid (HA) using 4-methylumbelliferone (4-MU). (H and I) Gross images of control (left) versus 4-MU (right) digits at 14DPA (top panels) or 28DPA (bottom panels). The black dashed line indicates the border of the nails, whose lengths (top) and areas (bottom) were quantified in (I). Scale bars = 100 µm. n = 5 mice per condition. (J-L) Microcomputed tomography analysis of skeletal morphologies of control and 4-MU digits at 14DPA (J) and 28DPA (K). Scale bars = 500 µm. The bottom panels (J’, J”, K’, and K”) show magnified views of the green boxes. The colored arrows indicate key events during the wound healing process, and the white dashed lines indicate where the bone length measurements were taken. Scale bars = 250 µm. The bone volume and length of the 28DPA digits were quantified in (L). n = 5 mice per condition. (M-O) Immunofluorescence images of 14DPA control and 4-MU digits stained for the blastema marker ARSI (M; white), Ki67 (N; magenta) and PDGFRα (N; white). All sections were counterstained with Hoechst (blue). The green dashed line indicates the border of the P3 bone. Scale bars = 250 µm. Insets show regions of the blastema (M’, N’) and wound (M”, N”) tissues at higher magnification. Scale bars = 50 µm. The number of Ki67^+^ cells per field of view (FOV) was quantified in (O). n = 3 mice per condition. (P) Immunofluorescence images of 28DPA control and 4-MU digits stained for the osteoblast marker SP7 (magenta), with SHG detection of collagen I (white). The yellow dashed line indicates the border of the P3 bone. Scale bars = 100 µm. Insets show higher magnification views of the osteogenic front of the control (P’) and 4-MU (P”) digits. Orange arrowheads indicate nuclear SP7 staining. Scale bars = 20 µm. Data are shown as mean ± SEM. Statistical significance was determined by two-tailed unpaired student’s *t*-test. *p < 0.05, **p<0.01, ***p<0.001, ns = not significant.

To examine the effects of continuous HA depletion, we incorporated 4-methylumbelliferone (4-MU) into the mouse diet prior to regenerative amputations (Figure 3G). 4-MU depletes UDP-glucuronic acid, one of two substrates required for HA synthesis^91^. With reduction of extracellular HA (Figures S3B and S3C), 4-MU digits at 14DPA were markedly reduced in length and area (Figures 3H and 3I). By 28DPA, control digits regenerated, but 4-MU digits remained significantly smaller (Figures 3H and 3I). To further characterize regeneration deficits observed with HA depletion, we analyzed digit bone structure using microcomputed tomography. Typically, 14DPA regenerating digits will have completed histolysis—the expulsion of a distal bone fragment resulting in bone shortening—prior to blastema formation and bone elongation^92,93^. Indeed, we found that these features were present in our control digits (Figure 3J). Conversely, 4-MU delayed histolysis-induced bone shortening (Figure 3J), leading to no difference in bone volume, surface area, or length between conditions (Figure S3D). By 28DPA, when regeneration is expected to be mostly completed^92^, control digits appropriately elongated (Figures 3K, 3L, and S3E), while 4-MU digit bone volume, surface area, and length remained markedly smaller (Figures 3K, 3L, and S3E).

We then examined the state of the blastema in 4-MU digits, hypothesizing that the digits’ defects were a result of compromised blastema formation. Our previous work identified that the blastema uniquely expressed *Arsi*^8^, which we presently confirmed (Figure 3M). However, with 4-MU, wounded digits downregulated their expression of ARSI (Figure 3M). Another defining feature of the blastema is the prevalence of proliferative cells^93^. Expectedly, we found that the blastema had high numbers of Ki67^+^ proliferating cells, which was halved with 4-MU treatment (Figures 3N and 3O). Lastly, blastema fate involves differentiation into mature cell types of the regenerated tissue^10^. To assess the differentiation of skeletal cells at 28DPA, we used SP7 (OSX)^94^ to label osteoblasts. The regenerated digit contained a robust population of SP7^+^ osteoblasts, while 4-MU reduced osteoblast numbers (Figures 3P and S3F). Additionally, fibrosis-like collagen was deposited in 4-MU digits (Figures 3P and S3G), further indicating a switch to a more non-regenerative ECM. Altogether, depletion of HA interfered with digit restoration after distal tip amputation, likely due to a failure of blastema formation and differentiation. Our analysis, taken together, strongly suggested an indispensable role for HA in digit tip regeneration.

### Collagen assembly prevails in the absence of hyaluronic acid

Given that hyaluronic acid (HA) and collagen were inversely correlated in wounded digits, and HA depletion elicited fibrosis-like ECM, we asked whether HA matrices abrogate fibrotic collagen assembly. To answer this question, we first compared the collagen architecture of hyaluronidase-treated versus control digits at 14DPA (Figure 4A). Hyaluronidase treatment increased collagen content, as well as fiber alignment, width, and length, indicating fibrotic tissue progression (Figures 4B-4D, S4A, and S4B). We observed similar fibrotic architectural features after HA depletion using 4-MU (Figures 4E-4H, S4C, and S4D). Overall, disruption of HA reproduced collagen networks consistent with fibrosis.

**Figure 4.**
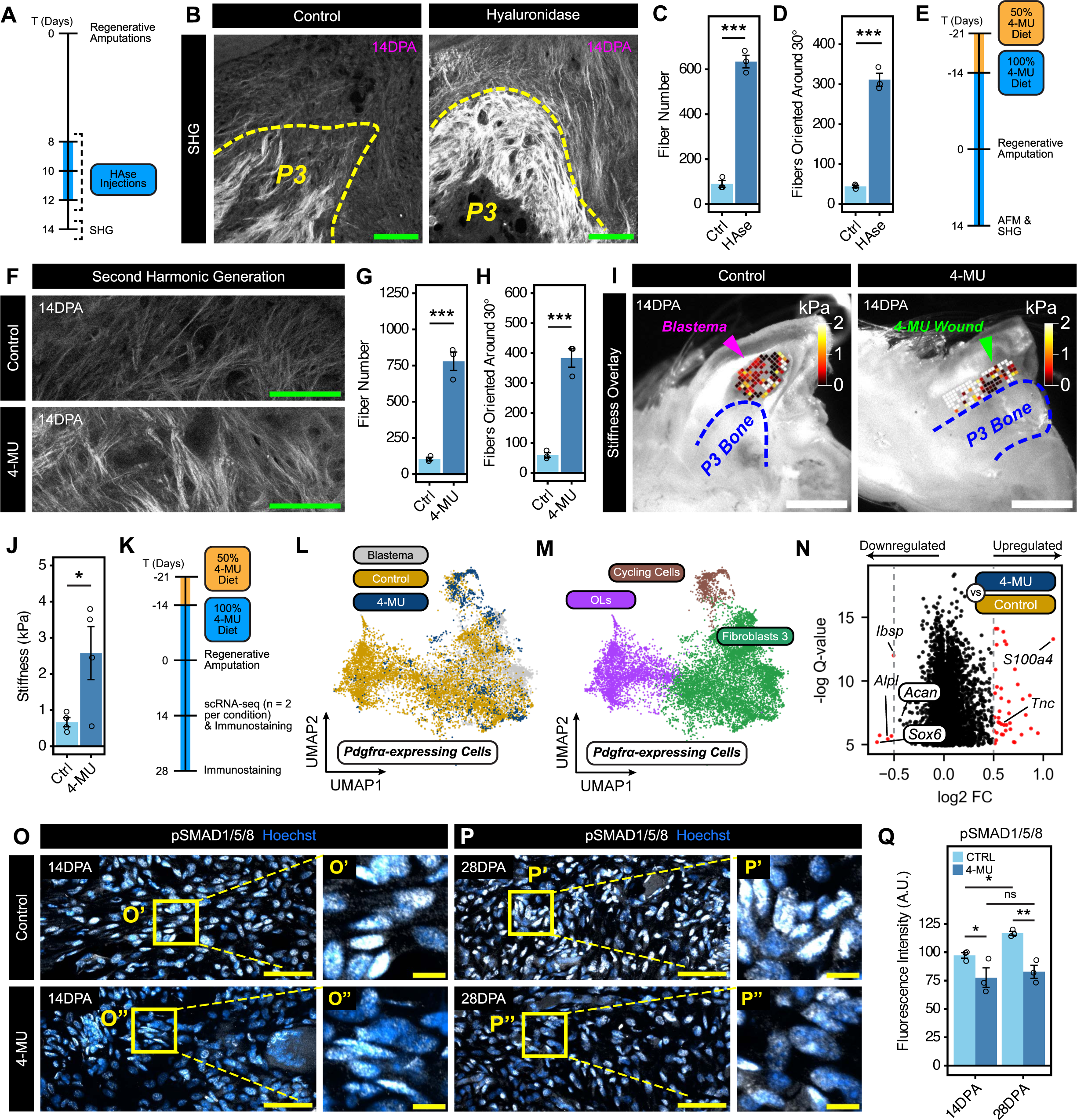
Hyaluronic acid regulates tissue mechanics and osteogenic differentiation. (A) Schematic of the experimental design testing the impact of hyaluronidase (HAse) degradation of hyaluronic acid on collagen fibrillogenesis. SHG = second harmonic generation microscopy. (B-D) Second harmonic generation (SHG) microscopy images of 14 days post-amputation (DPA) control and HAse digits highlighting collagen I fibers. The yellow dashed line indicates the border of the third phalanx (P3) bone. Scale bars = 100 µm. Fiber number (C) and orientation (D) were quantified using CT-FIRE. n = 3 mice per condition. (E) Schematic of the experimental design testing the impact of 4-MU depletion of hyaluronic acid on collagen and tissue mechanics. AFM = atomic force microscopy, SHG = second harmonic generation microscopy. (F-H) Second harmonic generation microscopy images of 14DPA control and 4-MU digits highlighting collagen I fibers. Scale bars = 100 µm. Fiber number (G) and orientation (H) were quantified using CT-FIRE. n = 3 mice per condition. (I and J) Atomic force microscopy stiffness maps of the 14DPA blastema (magenta arrowhead) and 4-MU wound (green arrowhead). The blue dashed line indicates the border of the P3 bone. Scale bars = 500 µm. Average stiffnesses were quantified in (J), which were determined by applying the Hertz model. n = 4 mice per condition. (K) Schematic of single-cell RNA sequencing and immunostaining experiments to determine the impact of 4-MU depletion of hyaluronic acid on cellular behavior. (L-M) UMAP of 8,995 *Pdgfrα*-expressing cells from the 4-MU and control datasets. Given the reduced biological replicates in the 4-MU experiment, prior 14DPA blastema datasets (gray) were integrated only to strengthen dimensionality reduction plotting; all downstream analyses were performed comparing 4-MU versus control samples only. In (M), three distinct cell sub-types were identified. They included an osteo-lineage (OL, purple), Fibroblast 3 (green), and cycling cell (brown) clusters. n = 2 replicates per condition, 6 mice pooled per replicate. (N) Volcano plot of differentially expressed genes in 4-MU versus control *Pdgfrα*-expressing cells, determined by the two-part generalized linear model MAST. Genes of interest have been annotated. (O-Q) Immunofluorescence images of 14DPA (O) and 28DPA (P) control (top panels) and 4-MU (bottom panels) digits stained for pSMAD1/5/8 (white) and Hoechst (blue). Scale bars = 100 µm. The right-hand panels show the yellow boxed regions of the blastema (O’ and P’) and 4-MU wound (O” and P”) tissues at higher magnification. Scale bars = 10 µm. Quantification of the average pSMAD1/5/8 fluorescence intensities in arbitrary units (A.U.) is shown in (Q). n = 3 mice per condition. Data are shown as mean ± SEM. Statistical significance was determined by two-tailed unpaired student’s *t*-test or two-way ANOVA with Tukey’s multiple comparisons test. *p < 0.05, **p<0.01, ***p<0.001, ns = not significant.

### HA and collagen mediate tissue mechanics in the regenerating blastema

If collagen networks develop in the absence of robust HA matrices, we hypothesized that HA depletion would create a mechanical environment resembling that of a non-regenerative wound. To test this idea, we perturbed HA using 4-MU prior to regenerative amputations and performed stiffness and fluidity measurements using atomic force microscopy (Figure 4E). We found that 4-MU digits were more than three-fold stiffer compared to controls (Figures 4I and 4J). 4-MU also diminished the tissue’s fluidity by half (Figures S4E and S4F). Overall, the viscoelasticity of 4-MU digits closely mirrored that of non-regenerative wounds (Figures 1D and S1G). These perturbation experiments demonstrated that HA-collagen interactions impact the mechanical microenvironment of wounded digits. We speculate that a high HA:collagen ratio imparts the blastema with its soft, fluid properties, while the converse imbues the non-regenerative tissue with high stiffness and low fluidity.

### Depleting hyaluronic acid suppresses fibroblast expansion and osteogenic differentiation

To probe how tissue mechanics arising from HA depletion affect cell behavior, we performed single-cell analysis of post-amputated P3 digits after 4-MU treatment versus control (Figure 4K). Unsupervised clustering and integration of the datasets resulted in nine major cell types (Figure S4G and S4H). Hyaluronic acid depletion demonstrably reduced the relative proportion of fibroblasts (Figure S4I), indicating a suppression of the typical injury-induced cell expansion. Analyzing the *Pdgfrα*-expressing cells, we showed a diminished OL proportion after 4-MU treatment (Figures 4L, 4M, S4J, and S4K). Comparing all *Pdgfrα*-expressing cells between conditions underscored the downregulation of OL-related genes *Ibsp* and *Alpl* with HA depletion, as well as *Sox6* and its target gene *Acan* (Figure 4N). These results highlighted a disturbance in osteogenic differentiation and the downregulation of cartilage-specific matrix elements. Interestingly, 4-MU fibroblasts upregulated their expression of *S100a4* and *Tnc*, both of which are associated with increasing ECM stiffness^95,96^. Altogether, OLs not only help establish the ECM environment in regeneration but are also acutely sensitive to HA perturbations and changes in tissue mechanics.

### Hyaluronic acid regulates BMP signaling

Given the digit’s skeletal defects and disruption in osteogenic differentiation with HA depletion, we explored the relationship between HA, tissue mechanics, and the bone morphogenic protein (BMP) pathway. We assayed the primary effector of BMP signaling, pSMAD/1/5/8^97^, after HA depletion with 4-MU *in situ*. pSmad1/5/8 levels were elevated and increased as regeneration progressed in control digits (Figures 4O-4Q). In contrast, 4-MU persistently suppressed pSMAD1/5/8 levels in the wounded tissue (Figures 4O-4Q). These initial findings suggested that HA and possibly tissue mechanics regulate BMP signaling.

### Responsiveness to BMP ligands is increased on soft substrates

To test the possibility that tissue mechanics mediate BMP signaling, we cultured fibroblasts on soft and stiff hydrogels and administered a pulse of BMP-7—a ligand present in the blastema during digit regeneration^16^—or medium without BMP-7 as control (Figure 5A). We observed the highest nuclear pSMAD1/5/8 signal in stimulated fibroblasts on soft substrates (Figures 5B, 5C, and S5A). Immunoblotting further corroborated these stiffness-dependent patterns in pSMAD1/5/8 levels (Figures S5B and S5C). We additionally profiled changes in the gene expression of *Inhibitors of DNA Binding/Differentiation* (*Id*), to assess early downstream effectors of pSMADs^84^. *Id1* expression uniquely responded to substrate stiffness, with fibroblasts on soft substrates upregulating *Id1* the highest (Figures 5D and S5D). We corroborated similar trends among freshly isolated cells from the second and third phalanx regions, which upregulated their expression of *Id1* most when stimulated under soft conditions (Figure S5E and S5F). Collectively, our data strongly indicated that substrate stiffness tunes BMP signal transduction.

**Figure 5.**
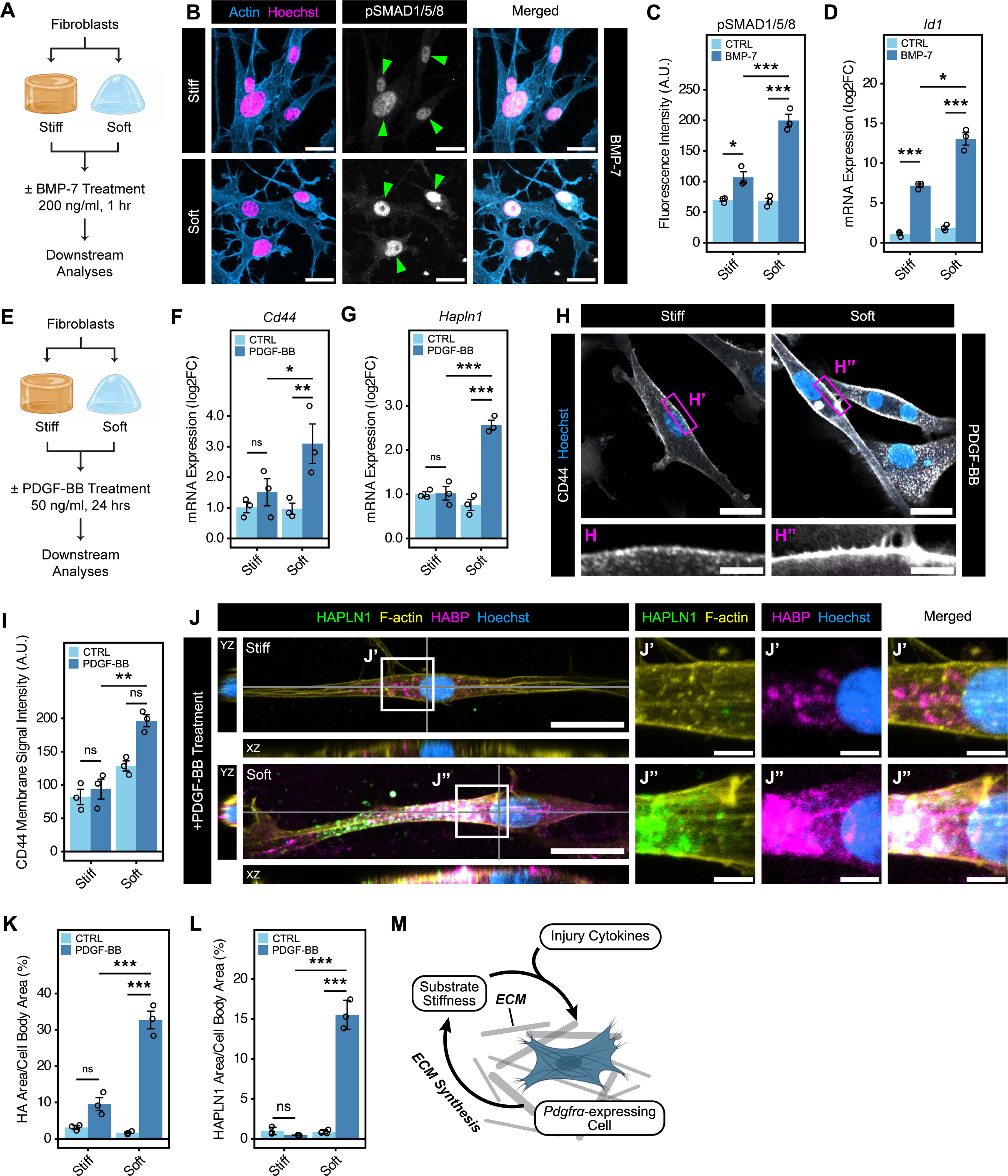
Substrate stiffness tunes BMP signaling and synthesis of pro-regenerative ECM. (A) Schematic of the experimental design to test the effects of substrate stiffness on BMP signaling. (B and C) Immunofluorescence images of BMP-7 treated fibroblasts cultured on stiff (50 kPa; upper panels) and soft (0.7 kPa; lower panels) hydrogels and stained for pSMAD1/5/8 (white), F-actin (blue), and Hoechst (magenta). Green arrowheads indicate nuclear pSMAD1/5/8 localization. Scale bars = 20 µm. The fluorescence intensities, presented as arbitrary units (A.U.), of pSMAD1/5/8 in cells cultured with, or as controls without, BMP-7 were quantified in (C). n = 3 independent experiments. (D) qPCR analysis of *Id1* gene expression in fibroblasts cultured on stiff (50 kPa) or soft (0.7 kPa) hydrogels, with or without BMP-7. n = 3 independent experiments. (E) Schematic of the experimental design to test the effects of substrate stiffness on PDGF-BB signaling. (F) qPCR analysis of *Cd44* gene expression in fibroblasts cultured on stiff (50 kPa) or soft (0.7 kPa) hydrogels, with or without PDGF-BB. n = 3 independent experiments. (G) qPCR analysis of *Hapln1* gene expression in fibroblasts cultured on stiff (50 kPa) or soft (0.7 kPa) hydrogels, with or without PDGF-BB. n = 3 independent experiments. (H and I) Immunofluorescence images of PDGF-BB treated fibroblasts cultured on stiff (50 kPa) or soft (0.7 kPa) hydrogels and stained for CD44 (white) and Hoechst (blue). Scale bars = 25 µm. The bottom panels show regions of the cell membrane (magenta boxes; G’ and G”) at higher magnification. Scale bars = 10 µm. The fluorescence intensities, presented as arbitrary units (A.U.), of CD44 in cells cultured with, or as controls without, PDGF-BB were quantified in (H). n = 3 independent experiments. (J-L) Immunofluorescence images of PDGF-BB treated fibroblasts cultured on stiff (50 kPa) or soft (0.7 kPa) hydrogels and stained for HABP (magenta), HAPLN1 (green), F-actin (yellow), and Hoechst (blue). XZ and YZ images are orthogonal views at the plane of the gray solid lines. Scale bars = 25 µm. The right-hand panels show the gray boxed regions (J’ and J”) at higher magnifications in split images. Scale bars = 5 µm. The percentage hyaluronic acid (K) and HAPLN1 (L) coverage in treated fibroblasts, with or without PDGF-BB, were quantified. n = 3 independent experiments. (M) Working model of *Pdgfrα*-expressing cell-stiffness feedback mechanisms that involve ECM synthesis and injury cytokines. Data are shown as mean ± SEM. Statistical significance was determined by two-way ANOVA with Tukey’s multiple comparisons test. qPCR data were normalized to Stiff-Control conditions and shown as log2FC, with statistical analyses performed on -11CT values. *p < 0.05, **p<0.01, ***p<0.001, ns = not significant.

### Soft substrates enhance the synthesis of pro-regenerative ECM

Since stiff mechanical cues enhance non-regenerative ECM synthesis (Figures 1N and 1O), we investigated whether a soft environment augments the production of “pro-regenerative” ECM. We plated fibroblasts on soft and stiff hydrogels and used Platelet-Derived Growth Factor-BB (PDGF-BB)—an injury signal with potent HA-synthesizing activity^98^—to test how substrate stiffness influences HA formation (Figure 5E). In addition to *Hapln1* and *Acan*, we considered factors that facilitate the assembly of HA networks. These factors include the three isoforms of hyaluronan synthases—*Has1*, *Has2*, and *Has3*—as well as *Cd44*, which tethers HA to the cell surface^99^. With PDGF-BB stimulation, fibroblasts upregulated *Has1-3*, but substrate stiffness had little effect on expression levels (Figure S5G). Under these experimental conditions, *Acan* was not responsive to either stimulation or substrate stiffness (Figure S5H).

In contrast, fibroblasts cultured on soft gels with PDGF-BB treatment significantly upregulated *Hapln1* and *Cd44* expression (Figures 5F and 5G). Furthermore, CD44 exhibited greater localization to the cell membrane (Figures 5H, 5I, and S5I). Reasoning that CD44 and HAPLN1 tether and confer structural integrity to HA networks, we asked whether fibroblasts assemble pericellular HA networks more robustly under soft conditions. After PDGF-BB treatment, fibroblasts cultured on soft substrates produced dense aggregates of HA and HAPLN1 at the cell-surface, with significantly greater surface coverage by both (Figures 5J-5L and S5J). Altogether, efficiency in assembling pericellular HA was stiffness-dependent, and our findings suggested that the tethering factors CD44 and HAPLN1 play more influential roles in HA network formation than HA synthases. Moreover, the softness imbued by HA may initiate positive feedback mechanisms to enhance the production of HA networks themselves, in contrast to stiff substrate-induced collagen fibrillogenesis (Figure 5M).

### HAPLN1 promotes the accumulation of HA and mediates collagen fibrillogenesis

Next, we investigated whether HAPLN1 itself promotes pericellular HA accumulation. Not only was HAPLN1 abundant alongside HA in the blastema (Figure 2J), but blastema-like substrate softness also propagated the synthesis of both (Figures 5I-5L). Analysis of uninjured digits further solidified the link between HAPLN1 and HA, whereby the P3 region contained wide swaths of HA-rich regions, which co-localized with HAPLN1 (Figure 6A). In contrast, this co-localization appeared only in specific P2 regions, such as the periosteum (Figure 6A). Regional differences in HAPLN1 were corroborated by gene expression analysis of P2 and P3 cells, in which P3 cells expressed higher levels of *Hapln1* (Figures 6B and 6C). These findings showcased strong associations between HAPLN1 and HA in injury and homeostasis, leading us to conclude that HAPLN1 is a key factor mediating the retention or stabilization of HA. Furthermore, since our findings indicate that the regulation of HAPLN1 was mechanosensitive, we subsequently explored its influence on HA and regeneration.

**Figure 6.**
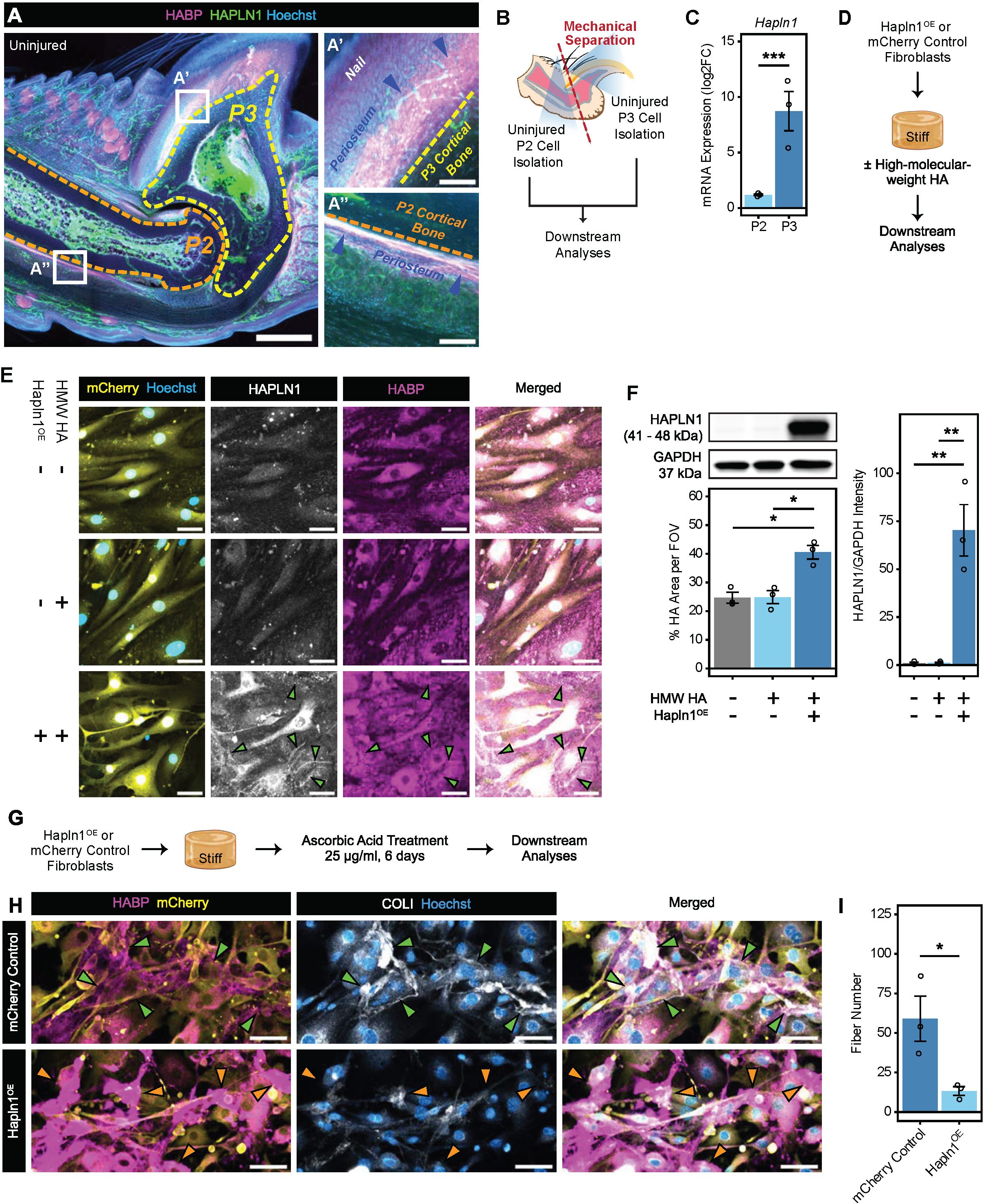
HAPLN1-mediated hyaluronic acid aggregation remodels collagen. (A) Immunofluorescence images of uninjured digits stained for HABP (magenta), HAPLN1 (green), and Hoechst (blue). The yellow dashed line indicates the border of the third phalanx (P3) bone, and the orange dashed line the border of the second phalanx (P2) bone. Scale bar = 300 µm. The right-hand panels (A’ and A”) show the white boxed regions of the P3 and P2 periosteum (blue arrowheads) at higher magnification. Scale bars = 50 µm. (B) Schematic of the experimental design testing the expression of *Hapln1* in uninjured P2 and P3 cells. (C) qPCR analysis of *Hapln1* expression in uninjured P2 and P3 cells. n = 3 independent experiments. (D) Schematic of the experimental design testing the impact of *Hapln1* overexpression (Hapln1^OE^) on pericellular hyaluronic acid in a stiff, fibrosis-like mechanical environment, with or without high-molecular-weight hyaluronic acid (HA). (E) Immunofluorescence images of mCherry Control and Hapln1^OE^ fibroblasts cultured on stiff (50 kPa) hydrogels, with or without high-molecular-weight (HMW) hyaluronic acid (HA) and stained for mCherry (yellow), HAPLN1 (white), HABP (magenta), and Hoechst (blue). The green arrowheads point towards bright regions of extracellular HAPLN1, co-localizing with aggregates of HA. Scale bars = 50 µm. (F) Immunoblotting for HAPLN1 with GAPDH as the loading control in mCherry Control and Hapln1^OE^ fibroblasts cultured on stiff (50 kPa) hydrogels, with or without HMW HA. Corresponding HA coverage per field of view (FOV) was quantified, related to Figure 6E. Quantification of HAPLN1 integrated band intensity normalized to that of GAPDH. n = 3 independent experiments. (G) Schematic of the experimental design testing the impact of *Hapln1* overexpression (Hapln1^OE^) on collagen fibrillogenesis, induced by ascorbic acid. (H and I) Immunofluorescence images of mCherry Control and Hapln1^OE^ fibroblasts cultured on stiff (50 kPa) hydrogels with ascorbic acid and stained for HABP (magenta), mCherry (yellow), COLI (white), and Hoechst (blue). The green arrowheads highlight dim regions of extracellular HA, co-localizing with bright collagen fibers. The orange arrowheads highlight bright regions of HA, co-localizing with dim collagen fibers. Scale bars = 50 µm. The number of collagen fibers per field of view was quantified in (I). n = 3 independent experiments. Data are shown as mean ± SEM. Statistical significance was determined by two-tailed unpaired student’s *t*-test, three-way ANOVA with Tukey’s multiple comparisons test, or one-way ANOVA with Tukey’s multiple comparisons test. qPCR data were normalized to P2 cells and shown as log2FC, with statistical analyses performed on -11CT values. *p < 0.05, **p<0.01, ***p<0.001, ns = not significant.

We then tested whether HAPLN1 influences the formation of HA networks by overexpressing *Hapln1* (Hapln1^OE^) or a scramble sequence as controls (mCherry Controls) in fibroblasts (Figures S6A-S6C). We cultured transduced fibroblasts on stiff hydrogels to simulate the fibrotic mechanical environment, with or without the presence of high-molecular-weight HA in the culture medium to encourage pericellular HA accumulation (Figure 6D). Despite HA supplementation, mCherry Control fibroblasts exhibited sparse and limited distribution of HA across their cell surface (Figures 6E and 6F). Meanwhile, Hapln1^OE^ fibroblasts synthesized large quantities of HAPLN1 that corresponded with a significant increase in pericellular HA (Figures 6E and 6F). Thus, despite exposure to mechanical stress, Hapln1^OE^ promoted the deposition of HA, signifying HAPLN1’s potential to promote regenerative ECM in a non-regenerative context.

Since depletion of HA increased matrix stiffness (Figure 4J), partly through greater collagen deposition, and stiff substrates upregulated fibroblasts’ synthesis of collagen (Figure 1N), we tested whether stabilizing HA networks using HAPLN1 could inhibit stiffness-induced collagen fibrillogenesis. To mimic the fibrogenic milieu, we cultured transduced fibroblasts on stiff hydrogels and used ascorbic acid to induce collagen fibrillogenesis (Figure 6G). mCherry Control fibroblasts exhibited low pericellular HA and upregulated synthesis of collagen fibrils (Figures 6H and 6I). By contrast, Hapln1^OE^ fibroblasts accumulated a robust network of pericellular HA that coincided with fewer and shorter collagen fibrils (Figures 6H, 6I, S6D, and S6E). Altogether, since HA restrains collagen fibrillogenesis, our data suggested that HAPLN1-augmented HA matrix may confer softening of the tissue environment.

### HAPLN1 promotes restorative repair after non-regenerative digit amputations

Since regeneration requires HA, and HAPLN1 fosters the formation of HA networks, we hypothesized that overexpressing *Hapln1* in non-regenerative wounds would initiate restorative repair. To test this hypothesis, we performed non-regenerative amputations on adult immunocompromised mice and transplanted mCherry Control fibroblasts or fibroblasts overexpressing Hapln1 (Hapln1^OE^) into the digit tip (Figure 7A). Using microcomputed tomography, we showed enhanced bone repair in digits injected with Hapln1^OE^ fibroblasts, including greater bone regrowth and elongation (Figures 7B-7D and S7B). Furthermore, at both 14- and 28DPA, elevated HAPLN1 promoted HA accumulation (Figures 7E, 7F, and S7C). Imaging of collagen structures revealed that regenerating bone tissue in Hapln1^OE^ digits extended well beyond the initial amputation plane, in contrast to minimal new tissue formation in controls (Figures 7G and 7H). Additionally, Hapln1^OE^ fibroblasts decreased the presence of fibrosis-like collagen structures (Figures S7D-S7H). Interestingly, we also observed an incidence of nail regrowth with Hapln1^OE^ cell transplantation, which does not occur under non-regenerative circumstances (Figure S7A). Taken together, we demonstrated that HAPLN1 triggers restorative repair after non-regenerative amputations, likely by mediating the HA- and collagen-matrices.

**Figure 7.**
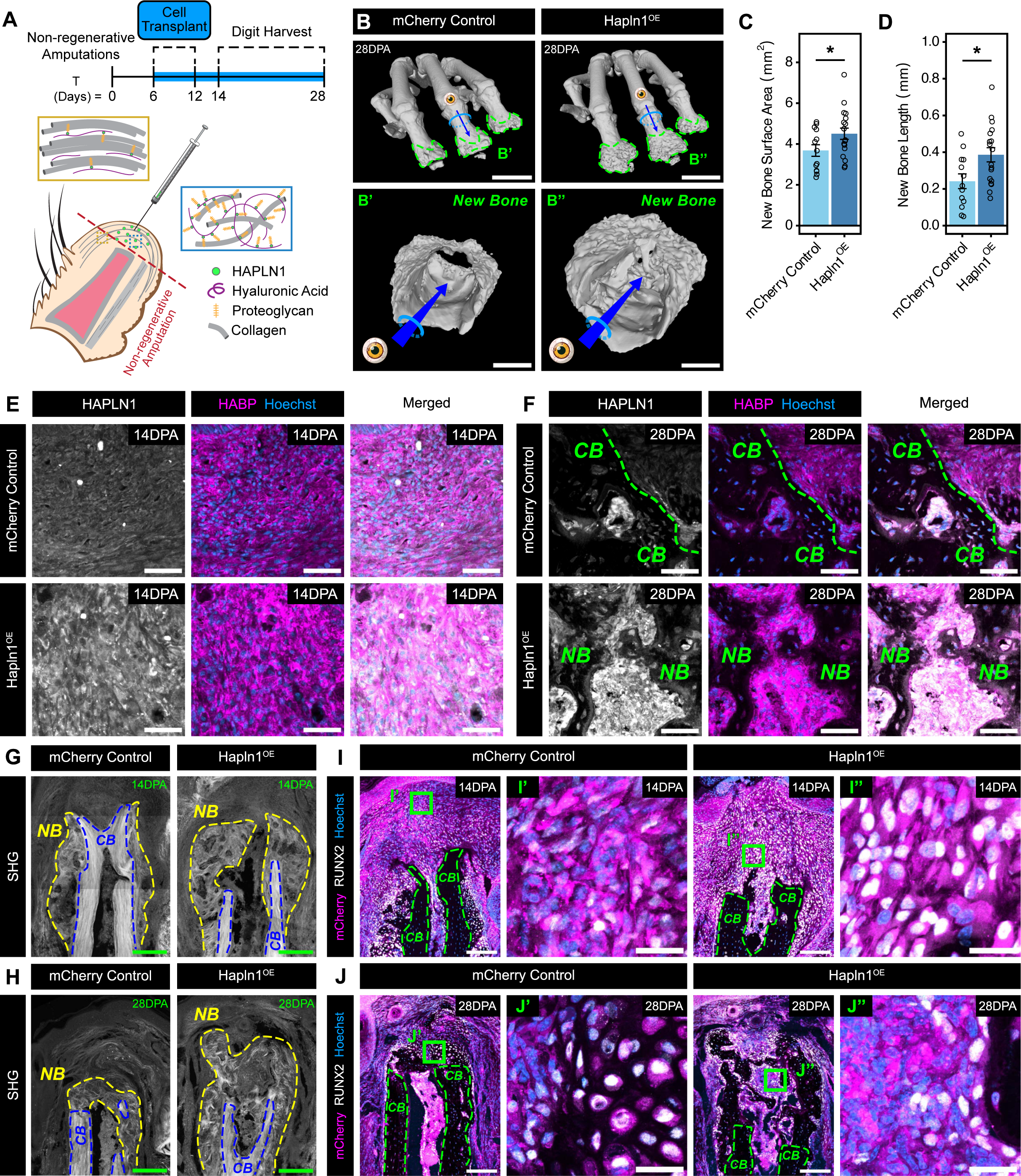
Hyaluronic acid initiates rescue of non-regenerative amputations. (A) Schematic of experimental design testing the transplantation of mCherry Control or *Hapln1*overexpressing fibroblasts in digits after non-regenerative amputations. (B-D) Microcomputed tomography analysis of 28DPA digits following transplantation of mCherry Control or *Hapln1* overexpressing (Hapln1^OE^) fibroblasts. The green dashed line indicates the border of new bone formation. Scale bars = 1 mm. The bottom panels show examples of new bone formation (B’ and B”) at higher magnification. Scale bars = 200 µm. The bone surface area (C) and length (D) of the 28DPA digits were quantified. n = 18 digits per condition. (E and F) Immunofluorescence images of 14DPA (E) and 28DPA (F) digits after transplantation of mCherry Control or Hapln1^OE^ fibroblasts and stained for HAPLN1 (white), HABP (magenta), and Hoechst (blue). The green dashed line indicates the border of the pre-existing cortical bone (CB). Scale bars = 50 µm. NB = new bone. (G and H) Second harmonic generation (SHG) images of 14DPA (G) and 28DPA (H) digits after transplantation of mCherry Control or Hapln1^OE^ fibroblasts. The blue dashed line indicates the border of the pre-existing cortical bone (CB), and the yellow dashed line indicates the border of new bone (NB) formation. Scale bars = 200 µm. (I and J) Immunofluorescence images of 14DPA (I) and 28DPA (J) digits after transplantation of mCherry Control or Hapln1^OE^ fibroblasts and stained for mCherry (magenta), RUNX2 (white), and Hoechst (blue). The pre-existing cortical bone (CB) is outlined by the green hashed lines. Scale bars = 150 µm. The right-hand panels show the green boxed regions (I’, I”, J’, and J”) at higher magnification. Scale bars = 25 µm. Data are shown as mean ± SEM. Statistical significance was determined by two-tailed unpaired student’s *t*-test. *p < 0.05, **p<0.01, ***p<0.001, ns = not significant.

Lastly, we explored how ECM modulation by HAPLN1 affects cellular activity. Since *Sox9* was elevated in regeneration (Figure 2C), we assayed SOX9 distribution as a marker for early osteogenesis^100^. Digits with mCherry Control fibroblasts exhibited low SOX9^+^ cell numbers, while clusters of SOX9^+^ cells localized distally from the plane of amputation in digits with Hapln1^OE^ fibroblasts (Figure S7I). We also assayed for more differentiated RUNX2^+^ OL cells. In digits transplanted with mCherry Control fibroblasts, we observed RUNX2^+^ osteo-lineage cells (OLs) only immediately adjacent to the pre-existing cortical bone at 14- and 28DPA (Figures 7I and 7J). However, Hapln1^OE^ fibroblasts increased the presence and distal localization of RUNX2^+^ OLs at both timepoints, consistent with bone regrowth (Figures 7I and 7J). These results showed that HAPLN1 markedly enhanced osteogenic differentiation. Altogether, we demonstrated a method of promoting cellular repair responses by targeting the ECM using HAPLN1.

## Discussion

In this work, we uncovered a mechanistic understanding of how the extracellular matrix (ECM) and emergent tissue mechanical properties drive regeneration in lieu of fibrosis. Specifically, our findings highlight three major conclusions. First, we identified ECM composition and tissue mechanics as key properties distinguishing non-regenerating wounds from the blastema, driven in part by cellular differences. In the former, two distinct *Pdgfrα*-expressing fibroblasts predominate, promoting stiff collagen networks through the production of collagen-remodeling factors. In contrast, osteo-lineage cells (OLs) contribute to the blastema’s soft ECM by synthesizing hyaluronic acid (HA) networks and associated components, such as ACAN and HAPLN1. Second, we demonstrated that HA is necessary for digit regeneration. Its perturbation prohibits restoration of amputated digits, while eliciting fibrotic ECM and tissue mechanics. Thirdly, our results support the conclusion that substrate mechanics reinforce wound healing outcomes by impacting cell behavior. For instance, stiff substrates propagate the synthesis of fibrotic ECM, whereas soft substrates promote regenerative ECM. Additionally, soft substrates enhance BMP signaling, which is central to skeletal regeneration^16,17,101^. Demonstrating partial digit restoration *in vivo* by promoting HA after non-regenerative amputations with HAPLN1 further supports these insights. Taken together, feedback processes among cells, the ECM, and tissue mechanics shape the wound healing outcome.

In two opposing wound healing conditions, our study shed light on the interplay of cell and ECM composition with tissue mechanics. Alongside recent recognition that fibroblasts are heterogenous, other studies have pinpointed cell sub-types that are responsible for fibrosis in different organ systems. For example, in wounded skin, expression of *Engrailed-1* in a subset of fibroblasts drives scar formation, and inhibiting its mechanical activation promoted skin regeneration without scarring^68,69^. Another study identified a distinct fibroblast sub-type that arises in injured hearts, and blocking their activation by immune cells reduced scar formation^102^. Similarly, our study highlighted specific *Pdgfrα*-expressing stromal cell populations that set non-regenerating wounds apart from the blastema and vice versa. We additionally provided a possible explanation for how these cell populations affect tissue mechanics by their production of collagen-remodeling factors. Fibroblast 1 and 2 cells produce high levels of TNX and THBS4 in non-regenerating wounds, both of which are positively associated with collagen remodeling^76,77,103–106^, with clear impact on tissue mechanics. Our work offers a framework for understanding how cell sub-types and their remodeling of the extracellular milieu govern the mechanical properties of tissues.

Through HA perturbation and rescue experiments in the context of digit tip amputations, our study strengthens the case that HA is critical for regeneration. Moreover, we posit that HA facilitates wound repair by modulating collagen fibrillogenesis and establishing tissue mechanics. While this phenomenon has been documented in fetal wounds^35–38^, our findings show that the HA-collagen relationship is conserved in injuries across developmental stages. However, the exact mechanism by which HA mediates collagen assembly remains to be elucidated. One possibility is that bulky pericellular HA matrices around osteo-lineage cells (OLs) regulate integrin accessibility to collagen ligands^107^. Another is HA’s physical interference of the self-assembly and organization of collagen polymers^108^, possibly through steric hindrance or by limiting diffusion of pro-collagen.

Lastly, we revealed that substrate mechanics directs cellular responses to soluble injury cues. Our evidence suggests that mechanical cues not only regulate fibrosis but also cellular capacity for regeneration. Prior studies have also linked mechanically soft environments to the maintenance of cellular function and even rejuvenation. One study found that elevated ECM stiffness compromises the expression of hematopoiesis-supportive factors by bone marrow stromal cells, and their culture in soft hydrogels partially rescued this deficit^45^. Another study showed that the brain stiffens with age, which diminishes the function of oligodendrocyte precursor cells^44^. Knockdown of the mechanosensitive ion channel Piezo1 reduced age-related changes to the oligodendrocyte precursor cells and improved their regenerative capacity after demyelinating injuries^44^. With the goal of modifying the non-regenerating wound’s ECM to resemble the blastema, we used HAPLN1 to successfully promote HA deposition. We also showed less fibrosis with HAPLN1, which aligns with a previous finding that HAPLN1 remodels collagen^109^. Previous studies have also shown that link proteins stabilize aggregates of HA, link proteins, and proteoglycans^88,110,111^. HAPLN1 possibly protects HA against fragmentation into low-molecular-weight species, which is commonly associated with inflammation and fibrosis^112,113^. Altogether, these results suggest that higher-order structural organization of HA—namely, complex formation with proteoglycans through link protein—is a very important player in tissue mechanics and its role in biological function. Moreover, our results raise the possibility that the ECM and tissue mechanics can be therapeutically targeted to promote desirable wound healing outcomes.

## Materials and Methods

## Resource availability

### Lead contact

Further information and requests for resources and reagents should be directed to and will be fulfilled by Mekayla A. Storer

## Materials availability

Animal strains used in this study are available from Charles River Laboratories.

## Experimental model details

### Mice

C57BL/6, CD1, and NOD SCID mice (strain codes: 027, 022, and 394, respectively) were obtained from Charles River Laboratories. The animals were maintained in a standard facility for rodents at the University of Cambridge. They were housed in vented cages with controlled temperature, humidity, and 12 hr light/dark cycles. Enrichment items provided in the cages included red tubes, shredded paper, and a loft. Mice were fed autoclaved maintenance diet (SAFE R105) and water *ad libitum*. Animals were randomly divided into control and treatment groups, and both male and female mice were used. This research was regulated under the Animals (Scientific Procedures) Act 1986 Amendment Regulations 2012 following ethical review by the University of Cambridge Animal Welfare and Ethical Review Body (AWERB). All procedures were conducted strictly following the relevant protocols defined in Home Office Project License PP2274970.

## Method details

### Digit amputation model

Digit amputations were performed on 6- to 8-week-old mice as described previously^8,114^. Briefly, mice were orally administered meloxicam (0.5 mg/ml) for analgesia and anesthetized with isoflurane. Then, using sterile scalpels, the distal one-third of terminal phalanges 2, 3, and 4 were amputated to induce a regenerative response. For non-regenerative amputations, the distal one-third of second phalanges were amputated. During the entire length of the surgery, mice were kept on a heating pad to ensure maintenance of normal body temperature. Sterile gauze was placed over the wound for hemostasis, and once achieved, mice were returned to their cages and monitored every 15 minutes for any adverse effects to surgery. Over the next three days, the mice were carefully monitored and weighed to ensure appropriate recovery.

### Tissue preparation, immunostaining, and microscopy

Mouse tissues were harvested and fixed with 4% paraformaldehyde (Fisher Scientific, 11481745) at 4 °C overnight and decalcified using 0.5 M ethylenediaminetetraacetic (EDTA, Merck, 324504-500ML) acid pH 7.0 – 7.2 for 14 days. Tissues were cryo-protected overnight in 30% sucrose (Thermo Fisher Scientific, 419762500), embedded in optimal cutting temperature compound (Avantor, 361603E), and rapidly frozen on dry ice. These samples were stored at -80 °C until use.

For deep imaging of digits, samples were processed for immunohistochemistry (IHC) as previously described^115^. Briefly, tissues were sectioned to the mid-sagittal plane and placed in room temperature PBS to remove the OCT. Tissues were blocked overnight in PBS with 5% bovine serum albumin (BSA, Merck, 10735078001), 10% dimethylsulfoxide (DMSO, Merck, D8418-50ML), and 0.5% Triton X-100 (Merck, X100-5ML) in PBS (deep confocal blocking buffer) and protected from light. Primary antibodies were diluted in deep confocal blocking buffer and incubated with samples for 3 days. Samples were washed twice in PBS with 0.1% Triton X-100 (PBS-T) for 1 hr per wash, and the third wash was performed overnight. Secondary antibodies were also diluted in deep confocal blocking buffer and incubated for 3 days. Samples were washed as before, but additionally, when performing nuclear stains, Hoechst was added at 20 µg/ml (Biotechne, 5117/50) in the final 1-hr wash. On the final day, an alcohol series was performed by adding 70%, 95%, and 100% ethanol, each for 10 mins. Samples were either stored for at most 2 wks in 100% ethanol or cleared using benzyl alcohol:benzyl benzoate (Merck, 108006-2.5L and B6630-250ML, respectively) at a 1:2 ratio for 1 hr. Lastly, samples were mounted on to glass bottom dishes (Thermo Fisher Scientific, 150682) for imaging.

For thin-tissue IHC, samples were sectioned sagittally at 14 µm thickness using the Leica CM1950 cryostat and mounted on SuperFrost Plus Adhesion slides (Fischer Scientific, 10149870). Prior to staining, sections were outlined with ImmEdge Hydrophobic Barrier PAP Pen (Vector Laboratories, H-4000) and heated for 10 mins at 37 °C. Subsequently, slides were soaked in PBS with 0.5% Triton X-100 for 30 mins and blocked for 1 hr with 5% BSA in PBS-T (thin-tissue blocking buffer). If primary antibodies were from goat or sheep hosts, then 10% donkey serum (Merck, D9663-10ML) in PBS-T was used. Primary antibodies were diluted in thin-tissue blocking buffer, and slides were incubated with antibodies overnight at 4 °C. The next day, slides were washed 4 times in PBS-T, 5 mins each. Secondary antibodies were diluted in thin-tissue blocking buffer and applied to samples for 2 hrs at room temperature with Hoechst. Slides were washed 4 times in PBS-T and finally mounted in ProLong Gold Antifade Mountant (Thermo Fisher Scientific, P36930) and allowed to set overnight before sealing with nail polish. Samples were stored at 4 °C.

For visualization of HA using hyaluronic acid binding protein (HABP, Merck, 385911-50UG), sections were incubated with streptavidin-peroxidase from Streptomyces avidinii (Merck, S5512-.1MG), followed by chromogenic development using the Pierce DAB Substrate Kit, as per the manufacturer’s instructions (Thermo Fisher Scientific, 34002). For fluorescence microscopy, streptavidin secondaries were used.

For phosphorylated SMAD1/5/8 IHC, antigen retrieval (10 mM sodium citrate dihydrate (Merck, W302600-1KG-K) with 0.1% Triton X-100 at pH 6.0) was performed on cryo-sectioned, formalin-fixed tissues for 2 hrs at 70 °C in a water bath. Samples were rinsed with PBS and dried for 10 mins at 37 °C. Sections were outlined with a hydrophobic pen and dried for another 10 mins at 37 °C. All remaining steps were performed using IHC procedures described above, except PBS was replaced with Tris Buffered Saline (TBS) (10X, 200 mM Tris (Scientific Laboratory Supplies, 93350-100G), 1500 mM NaCl (Merck, S9888-500G), pH 7.6).

For immunocytochemistry, cells were fixed with 4% paraformaldehyde for 15 mins and permeabilized in 0.5% Triton X-100 in PBS for 30 mins. Except for pSMAD staining, which required TBS, staining procedures were the same as described above. For actin visualization, phalloidin (Proteintech, PF00001) staining was performed for 30 mins. To image, cells cultured on hydrogels were inverted onto glass bottom dishes.

For all IHC experiments, at least three digits were analyzed per condition, with each digit sourced from a separate animal. To visualize samples, the following confocal microscopes were used: Zeiss LSM 880 NLO, Zeiss LSM 980 Airyscan2 (Zeiss, Oberkochen, Germany), and the Leica Stellaris 8 (Leica, Wetzlar, Germany).

### Atomic force microscopy

Freshly isolated, post-amputated digits were cut mid-sagittally using a sharp scalpel and partially embedded in 4% agarose to stabilize the tissues on glass-bottom dishes. After the agarose had set, the samples were submerged in low-glucose DMEM without phenol red (Thermo Fisher Scientific, 11880036) and maintained at 34 °C. Atomic force microscopy (AFM) indentation measurements were performed using the setup previously described^116^. Briefly, the mounted digits were placed on the microscope stage, and an AxioZoom V16 stereomicroscope (Zeiss, Oberkochen, Germany) was used to acquire brightfield images. These images were used to designate a rectangular grid containing 2D spatial information upon which indentation measurements were performed. The exposed tissues were measured with force distance spectroscopy with probes consisting of 37.28 µm diameter polystyrene beads glued to Arrow-TL1 cantilevers (nominal spring constant = 0.01 N/m, NanoWorld, Arrow TL1). During measurements, the cantilever was approached at 20 µm/s until an indentation force of 10 nN was reached. This force was maintained for 3 seconds, thus enabling the acquisition of the deformation over time (‘creep’).

The reduced apparent elastic modulus 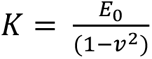 values were extracted by a custom-written MATLAB script applying the Hertz model^116^. A custom-written Python script implementing the Power Law Rheology (PLR) model (validated in Hecht et al^117^) was used to obtain fluidity values. *β* can have values ranging from 0 (for an elastic solid) to 1 (for a viscous fluid). For AFM experiments, a minimum of 4 separate digits were analyzed per condition, with each digit sourced from a separate animal.

### Digit cell and dermal fibroblast isolation and culture

To isolate uninjured or post-amputated P3 cells, the nails from C57BL/6 mouse digits were first removed to expose the underlying P3 bone and surrounding tissues. These tissues were digested in 0.250 mg/ml Liberase TH (Scientific Laboratory Supplies, 5401135001) in PBS (cell dissociation buffer) for 1 hr at 37 °C. The dissociated cells were treated with 20 U/ml DNase I (Merck, 4536282001) for 5 mins at 37 °C and passed through a 40 µm filter. Isolation of P2 cells was performed similarly as above. Briefly, P2 tissues were separated from P3 at the distal interphalangeal joint, and the skin was peeled from the P2 bone and minced. Together, the skin and the bone were incubated in cell dissociation buffer for 1 hr at 37 °C. Cells were maintained in standard fibroblast expansion medium, which contained 10% fetal bovine serum (FBS, Gibco, 10270106), 1% Penicillin/Streptomycin (Merck, P0781), and 10 ng/ml fibroblast growth factor 2 (FGF-2) (Peprotech, 100-18B) in low-glucose DMEM with pyruvate and HEPES (Thermo Fisher Scientific, 12320032). Cells were grown in 20% O2 and 5% CO2 in a 37 °C humidified incubator.

To isolate dermal fibroblasts, the back skin of CD1 mice at postnatal day 2 were dissected by using scissors to create a longitudinal incision across the length of the mouse, followed by manual separation of the skin. After scraping fat away using a sterile scalpel, the tissues were placed in 0.25% trypsin overnight at 4 °C. The next day, the dermis was separated from the epidermis using forceps and minced with scissors. The minced tissues were incubated in 1 mg/ml Collagenase P (Merck, 11213857001) and 2 mg/ml Dispase II (Merck, D4693-1G) and reconstituted in DMEM containing 2% FBS for 1 hr at 37 °C. DNA was digested using 20 U/ml DNase I for 5 mins at 37 °C. Fibroblast expansion medium was added to halt the digestion before passing the entire content of the tube through a 40 µm filter. Cells were spun for 10 mins at 300 x g and resuspended in fibroblast expansion medium.

### Single-cell isolation, capture, library construction, next generation sequencing, and alignment

For single-cell RNA sequencing (scRNA-seq), tissues from 14DPA non-regenerative (n = 2, 4 mice per replicate), 4-MU control (n = 2, 5 mice per replicate), and 4-MU (n = 2, 5 mice per replicate) digits were freshly dissected, and cells were isolated as described above, with an additional 30 µm filtering step to reduce tissue debris. The cells were resuspended at 372 cells/µl in 2% FBS in PBS. Libraries were prepared by using Chromium Next GEM Single Cell 3’mRNA v3.1 as per the company’s protocol and sequenced on a NovaSeq 6000 at the Cancer Research UK Institute (CRUK, Cambridge, UK). Raw 10X FASTQ files were aligned and quantified using the Cell Ranger Single-Cell Software Suite (v8.0.0, 10x Genomics). The mouse reference used was the mm10 reference genome refdata-gex-mm10-2020A, available at: https://cf.10xgenomics.com/supp/cell-exp/refdata-gex-mm10-2020-A.tar.gz.

### scRNA-seq data filtering, pre-processing, integration, clustering, and annotation

The 10X Genomics scRNA-seq datasets were processed in Python v3.10 using Scanpy v1.9.6. Publicly available mouse scRNA-seq blastema (n = 3) and non-regenerative datasets (n = 2) were acquired from Gene Expression Omnibus (GEO) under the accession numbers GSE135985^8^ and GSE143888^9^. Datasets that contained a high degree of cell-free (ambient) RNA molecules were cleaned with SoupX v1.6.2^118^ or CellBender v0.3.2^119^ before downstream pre-processing. Quality control for each dataset involved filtering based on fixed and batch-specific filtering thresholds. Fixed filtering thresholds included removing genes expressed in fewer than 3 cells and cells in which over 10% of the unique molecular identifiers arose from the mitochondrial genome. Batch-specific thresholds included removing cells beyond the upper and lower boundaries for number of genes and number of counts per cell. Doublets were identified using Scrublet v0.2.3^120^, with a threshold doublet score set at 0.2 to 0.25. The details of the filtering paraments are listed in Supplementary Table 1.

During batch pre-processing, size factors for each batch were calculated for data normalization using the computeSumFactors() in R v.4.3. The size factor normalized data were then subjected to logarithmization (scanpy.pp.log1p), and highly variable genes were detected (scanpy.pp.highly_variable_genes). Principal component analysis was performed by scanpy.pp.pca and the first 30 principal components were used to calculate neighbors (scanpy.pp.neighbors) and perform UMAP (scanpy.tl.umap) constructions. Leiden graph-based clustering (scanpy.tl.leiden) at default settings was used for broad cell type classification with manual annotation. For integration, the top 2,000 highly variable genes were sub-setted from the concatenated datasets. scVI v1.0.4 was used to integrate the datasets, with parameters set to n_latent = 30, n_layers = 2, and gene likelihood = “nb”. Training was conducted for a maximum of 800 epochs or stopped early based on elbo_validation. The latent representation obtained from the model was used for computing neighbors. The final clustering of cell populations was performed by selecting the most conservative resolution that 1) yielded distinct populations and appropriate heterogeneity based on the UMAPs and 2) retained high and differential expression of canonical cell type-specific markers. *Pdgfrα*-expressing clusters were sub-setted from the global integrated datasets for further analyses, and PCAs and UMAPs were re-calculated using scVI modeling and Leiden clustering to reveal sub-populations. The clustering resolution was determined as described above. Cell types were defined by finding marker genes with Scanpy (scanpy.tl.rank_genes_groups) using the Wilcoxon method and corrected by the Benjamini-Hochberg method.

### Proportional analysis, differential gene expression analysis, gene set enrichment analysis, and gene list scoring

Cell type proportion analyses were conducted with Propellor (Speckle v1.2.0)^121^, and p < 0.05 designated statistically significant differences between two groups. To account for technical variability across datasets, differential gene expression was performed using MAST v.1.8.2^122^, a two-part generalized linear model that factors in both the expression rate of each gene across cells and the mean gene expression. MAST analyses were used to compare the Fibroblast 1 and 2 clusters against all other cell types, as well as against OLs. For the 4-MU experiment, MAST analysis was performed on all *Pdgfrα*-expressing clusters between control and 4-MU datasets. Unless otherwise indicated, differentially expressed genes (DEGs) were determined as log2fold-change ≥ 0.5 and q-value < 0.05. The resulting DEGs were input for ClusterProfiler v4.10.1^123^ for gene ontology functional enrichment analysis. For complete Matrisome and collagen and proteoglycan scoring, ECM-related genes were taken from *mus musculus* complete matrisome list from the Matrisome Project^124^, and gene list scores were calculated using sc.tl.score_genes.

## Data and code availability

All raw scRNA-seq expression matrices from this project are available in the NCBI Gene Expression Omnibus under the accession number GSE274858. All script used for analysis and creating figures are available upon request.

### Fabrication of polyacrylamide gels of different stiffnesses

StemBond hydrogels were fabricated as previously described^82^. Briefly, support coverslips (VWR, 631-1581) were treated with 0.2 M sodium hydroxide (Merck, 567530-250GM) for 35 mins, cleaned, and functionalized with 3-(Trimethoxysilyl)propyl methacrylate (Merck, M6514-25ML) for 2 hrs. Top coverslips were treated with dichlorodimethylsilane (Merck, 440272-100ML) for 5 mins, polished with 100% ethanol, and cleaned. Hydrogel solutions were prepared according to Supplementary Table 2 with the inclusion of 6-acrylamidohexanoic acid (Tokyo Chemical Industry, A1896) and degassed in a vacuum chamber for 20 mins. To polymerize the hydrogels, TEMED and APS were added, and the hydrogels were sandwiched between the support and top coverslips for 35 mins. The hydrogels were hydrated overnight in 1% Penicillin-Streptomycin in PBS. The next day, top coverslips were detached from the hydrogels under sterile conditions and equilibrated with MES buffer (0.1 M MES hydrate (Merck, M2933-25G), 0.1 M NaCl, pH 6.1). Hydrogels were activated by a 30-min treatment with 0.2 M EDAC (Scientific Laboratory Supplies, 03450-25G) and 0.5 M NHS (Thermo Fisher Scientific, 157272500) in MES buffer with constant rocking. After, hydrogels were coated with poly-d-lysine hydrobromide (Merck, P0899-10MG) reconstituted in HEPES buffer (0.05 M HEPES (Merck H3375-25G), pH 8.5) at 100 µg/ml. Coating was performed on a rocker for 2 hrs at room temperature. Coating solution was aspirated, and the hydrogels were blocked with 0.5 M ethanolamine (Merck, E9508-100ML) in HEPES buffer. The hydrogels were thoroughly washed with several exchanges of PBS and stored at 4 °C for later use.

To test how substrate stiffness influences cell behavior, cells were cultured on stiff (50 kPa) and soft (0.7 kPa) hydrogels to mimic the mechanical microenvironment of the blastema and fibrotic tissue, respectively. Cells of no more than passage 4 were attached to the hydrogels for 24 hrs before treatment with PDGF-BB (50 ng/ml, Peprotech, 100-14B) or BMP-7 (200 ng/ml, Peprotech, 120-03P) in DMEM containing 1% FBS for experiments lasting less than or equal to 24 hrs. For longer experiments, DMEM with 10% FBS was used to maintain cell viability. To induce collagen synthesis and fibrillogenesis, 25 µg/ml 2-phospho-L-ascorbic acid trisodium salt (Merck, 49752-10G) was supplemented into the medium and replenished every other day until the experimental endpoint. For all hydrogel experiments, at least 3 independent experiments were performed.

### Hyaluronic acid extraction and enzyme-linked immunosorbent assay

To isolate hyaluronic acid (HA) from tissues, non-regenerating wounds and blastemas were micro-dissected, weighed, and incubated with digestion buffer containing 0.1 M Tris, 0.15 M NaCl, 0.01 M CaCl2 (Merck, 223506-25G), 5 mM deferoxamine mesylate (Merck, D9533-1G), and 0.5 mg/ml Proteinase K (Merck, P6556-5MG) at pH 8.3 for 2 hrs with vortexing every 30 mins. Samples were boiled at 100 °C for 20 mins to heat-inactivate Proteinase K and chilled on ice. 1 µl of Benzonase (Merck, E1014-5KU) was added to each sample and incubated for 1 hr to degrade DNA and RNA. Then, samples were centrifuged for 15 min at 21,000 x g at 4 °C, and the supernatant was collected into a separate tube. An equal volume of phenol:chloroform:isoamyl alcohol (Merck, 77617-100ML) was added, and after vortexing, the samples were centrifuged for 15 mins at 14,000 x g and 4 °C to separate the aqueous components from the other organics. This step was repeated using pure chloroform (Merck, 288306-100ML) to remove residual phenol from the aqueous phase. HA was precipitated using 100% ethanol and spun at 10,000 x g for 5 mins and washed with an additional volume of ethanol. The samples of extracted HA were resuspended in water. An ELISA to quantify the amount of HA in digit tissues was performed according to the manufacturer’s instructions (Biotechne, DHYAL0). Digits from four mice were pooled for a single biological replicate.

### *In vivo* knockdown of HA networks

For intermittent knockdown of HA, mice that underwent regenerative amputations were administered 0.4-1 U per digit of hyaluronidase from bovine testes (Merck, H3506-100MG) reconstituted in 0.1% BSA. Control mice received 0.1% BSA alone. Administration of substances was performed at 8-, 10-, and 12DPA using a 33G Hamilton syringes (Hamilton Company, 7635-01). Digits were harvested at 14DPA for processing and further analyses. Each digit was considered a biological replicate.

For continuous knockdown of HA, 4-methylumbelliferone (4-MU) was incorporated into chocolate-flavored chow (ssniff Spezialdiäten, Soest, Germany) at 50 g/kg, resulting in an approximate daily uptake of 250 mg of 4-MU per mouse. Control mice received the same chocolate-flavored chow without 4-MU. To acclimatize the mice to the modified diet and prevent weight loss, they were first fed maintenance chow:modified diet at a 50:50 ratio for the first seven days. Then, mice were maintained on the modified diet until the experimental endpoint. After acclimatization, mice underwent regenerative amputations, and tissues were collected at 14- and 28DPA for further analyses. Each mouse was considered a biological replicate.

### Reverse transcription quantitative polymerase chain reaction

To measure gene expression levels, cells were first lysed by direct addition of TRIzol (Thermo Fisher Scientific, 15596026) for 5 mins. Further RNA processing and purification was performed using Direct-zol RNA Microprep Kits (Zymo Research, R2060) according to the manufacturer’s instructions. Briefly, an equal volume of 100% ethanol to TRIzol was added to samples and spun in columns for two rounds of washing before elution in DNase/RNase-free water. Next, samples were treated with RQ1 RNase-free DNase (Promega, M6101) for 30 mins at 37 °C, followed by termination of the reaction using the supplied Stop Solution for 10 mins at 65 °C. RNA quality and concentration was measured using a Nanodrop Spectrophotometer (Thermo Fisher Scientific, Waltham, MA). Complementary DNA (cDNA) synthesis was carried out by first heating the RNA-primer mix containing 2.5 µM random hexamers (Thermo Fisher Scientific, N8080127), 0.5 mM dNTP (Thermo Fisher Scientific, 18427013), and 400 ng total RNA for 5 mins at 65 °C and chilling on ice for 1 min. Next, a reverse transcriptase mix containing 1x SuperScript IV, 5 mM DTT (Thermo Fisher Scientific, 18090010), and 2.0 U/µl RNaseOUT RNase Inhibitor (Thermo Fisher Scientific, 10777019) was added to each tube, which was incubated at 23 °C for 10 mins, 55 °C for 10 mins, and finally 80 °C for 10 mins. cDNA was diluted 1:200, and each PCR reaction contained 2 ng of cDNA, PowerTrack SYBR Green Master Mix (Thermo Fisher Scientific, A46109), and 800 nM of primer pairs. Reactions were carried out using the QuantStudio Real-Time PCR machine and software (Thermo Fisher Scientific, Waltham, MA). For all qPCR experiments, at least three independent experiments were performed. Statistical analyses were performed on −ΔΔ’(values, but expression graphs were depicted as fold change (2^$ΔΔ^()).

### Microcomputed tomography

To assess the skeletal morphology of mouse digits, 4% paraformaldehyde-fixed samples were scanned at either 14- or 28DPA using a Nikon XTEK 225 Micro CT Scanner (Nikon, Tokyo, Japan).

Scan settings were the following: 2.9-4.9 µm resolution, energy at 75-90 kV and 30-48 µA, zero filtration, 708-1000 ms exposure, 1080 projections, and two frames per projection. Image processing was done using CT Pro 3D and CT Agent (both Nikon, Tokyo, Japan). Scans were exported as .tiff stacks. The Materialise Mimics software (Materialise, Leuven, Belgium) was used to model samples in 3D and calculate volume and dimension measurements. For the 4-MU experiment, each mouse was considered a biological replicate, and for experiments involving direct injections of substances, each digit was considered a biological replicate.

### Western blotting

For immunoblotting, adherent cells on hydrogels were inverted onto Parafilm with RIPA buffer (Thermo Fisher Scientific, 89900) containing Halt Protease and Phosphatase Inhibitor Cocktail (Thermo Fisher Scientific, 78440). After 5 mins, a cell scraper was used to remove remaining adherent cells. The cell lysate was collected into a microcentrifuge tube and agitated for 30 mins at 4 °C. Then, the lysate was centrifuged for 5 mins at 14,000 x g and 4 °C. The supernatant was aspirated, and the total protein was quantified using the BCA method (Thermo Fisher Scientific, 23225). Samples were boiled in Laemmli sample buffer (5x, 250 mM Tris base, 5% SDS (Merck, L3771-25G), 50% glycerol (Merck, G7757-500ML), and 0.1% bromophenol blue (Merck, 114391-5G) with 2.5% 2-mercaptoethanol (Merck, M6250-10ML) at 100 °C for 5 mins, and 10 µg of total protein per lane, along with a Precision Plus Protein Kaleidoscope Prestained Protein Standards ladder (Bio-Rad Laboratories, 1610375), was separated by gel electrophoresis. Transfer of proteins to PVDF membranes was performed using the tank transfer method with Towbin buffer (10x, 0.25 M Tris base, 1.92 M glycine). The membranes were blocked with 5% BSA in TBS with 0.1% Tween-20 (Merck, P9416-50ML) (TBS-T) for 1 hr with constant agitation, followed by immunostaining with primary antibodies in blocking buffer overnight at 4 °C. Six 5-min washes with TBS-T were performed, followed by application of secondary antibodies for 1 hr in blocking buffer. After another six 5-min TBS-T washes, protein was detected using SuperSignal West Pico PLUS Chemiluminescent Substrate (Thermo Fisher Scientific, 34577) according to the manufacturer’s instructions. Visualization was performed using the G:Box chemi XRQ machine (Syngene, Cambridge, UK). Quantitative analysis of the immunoblots were performed using FIJI^125^. Densitometry analysis was performed by calculating background-subtracted integrated densities of pSMAD1/5/8 or HAPLN1 normalized to the loading control GAPDH. For all western blots, at least three independent experiments were performed.

### Overexpression of *Hapln1* in dermal fibroblasts

pLV[Exp]-EF1A-mHapln1-mCherry (Hapln1^OE^) and pLV[Exp]-EF1A-Scramble-mCherry (mCherry control) transfer plasmids were cloned and transformed in *Escherichia coli* (*E. coli*) by VectorBuilder (VectorBuilder, Chicago, IL). 3^rd^ generation lentivirus plasmids pMDLg/pRRE, pmD2.G, and pRSV-Rev (Addgene plasmids #12251, #12259, and #12253, respectively) were used to generate lentivirus containing the Hapln1^OE^ or mCherry Control transfer plasmid. HEK293T cells of passage less than 20 were seeded at a density of 4x10^6^ cells per 10-cm dish in high-glucose DMEM containing HEPES (Thermo Fisher Scientific, 10564011) and 10% FBS and allowed to attach overnight. The next day, fresh medium was supplied to the cells, which were subsequently transfected with the transfer, envelope, and packaging plasmids at a ratio of 4:2:1:1 by size (transfer:pMD2.G:pMDLg/pRRE:pRSV-Rev). The transfectant reagent:DNA complexes were made by combining *Trans*IT-VirusGEN (Mirus Bio, MIR 6704) with plasmids in Opti-MEM I (Thermo Fisher Scientific, 31985062) and incubating for 30 mins to form complexes. The *Trans*IT-VirusGEN:DNA complexes were added drop-wise to different areas of the dish. Lentivirus was collected at 48- and 72 hrs post-transfection and passed through a 0.45 µm filter. The lentivirus was then concentrated by ultracentrifugation at 25,000 RPM for 2 hrs at 4 °C. The supernatant was discarded, and the virus was resuspended in 100 µl PBS per 10-cm dish. Aliquots of lentivirus were then frozen at -80°C for later use.

Lentivirus multiplicity of infection and titer calculation was performed using a dilution series. The day prior to transduction, fibroblasts were plated at a density of 15,000 cells/cm^2^ in a 24-well plate. A serial dilution of lentivirus was added to the wells with the addition of 4 µg/ml of hexadimethrine bromide (Merck, H9268-10G). The plate was spun at 1,000 x g for 2 hrs at 32 °C, after which the medium was replaced with fresh fibroblast expansion medium. After 48 hrs, the percentage of reporter-positive cells was calculated. The volume of virus needed to achieve at least 95% infection efficiency was used to transduce fibroblasts for all experiments using the steps described above.

### Flow cytometry analysis

To confirm lentivirus transduction efficiency, infected cells were harvested and passed through a 40 µm filter. Then, cells were incubated with ready-to-use DAPI (Miltenyi Biotec, 130-111-570) for 5 mins in 2% FBS and 2.5 mM EDTA in PBS. Unstained fibroblasts, fibroblasts stained with DAPI, and mCherry-positive cells were used as single-stained controls for compensation and gating. Typically, 100,000 events were recorded for each sample. Data were acquired on a BD Fortessa flow cytometer (BD, Franklin Lakes, NJ), and data analysis was performed in the FlowJo v10.1 software (FlowJo, Ashland, OR). Three independent experiments were performed for quantification of transduction efficiency.

### *In vivo* transplantation of genetically modified dermal fibroblasts

Genetically modified fibroblasts were expanded, and cells of passage no later than 6 were used for transplantation. Briefly, detached cells were washed in PBS and resuspended at a concentration of 150,000 cells/µl in 33% EncapGel (Merck, 922412-1EA). NOD SCID mice, having undergone non-regenerative amputations, were administered 1 µl of cells or vehicle in the non-regenerative stump using a 33G Neuros Syringe at 6- and 12DPA. Mice digits were harvested at 14- and 28DPA for further processing. Each digit was considered a biological replicate.

### Quantification and statistical analysis

#### CT-FIRE analyses of second harmonic generation microscopy

To characterize collagen architecture in digits, second harmonic generation (SHG) microscopy with excitation wavelength of 920 nm was used to visualize collagen fibers overlying the P2 or P3 bone. Then, fibers were segmented in batch using CT-FIRE^126^ with the following parameters: threshold_im2: 65, s_xlinkbox: 5, and number of selected scales: 4. The specificity and sensitivity of the automatic segmentation were evaluated for each sample in overlaid images. CT-FIRE’s output consisted of each fiber’s width, length, number, and angle. To quantify the number of fibers oriented around 30°, a custom R script was written. Briefly, the most common fiber angle for a sample was designated as 0°, around which all other fiber angles were oriented. The total number of fibers oriented ± 30° were quantified and averaged for analysis between conditions. Two sections per digit were analyzed, and these counts were averaged and considered as a single biological replicate. A minimum of three separate animals were analyzed per condition, with each digit sourced from a separate animal. For experiments involving direct injections, each digit was considered a biological replicate. For collagen analysis of cells cultured on hydrogels, a minimum of three independent experiments were performed, and each hydrogel was imaged at 5 different locations, which were averaged as a single biological replicate. For collagen analysis of the Hapln1^OE^ experiment, three digits per condition were randomly chosen.

#### Quantification of percentage RUNX2^+^ cells after digit amputation

Using FIJI, the Hoechst channel was used to determine the total number of nuclei. The RUNX2 channel was used to quantify the total number of RUNX2^+^ nuclei. Percentage RUNX2-positivity was quantified as the number of RUNX2+ cells/total number of cells. Two sections per digit were analyzed, and these values were averaged and considered as a single biological replicate. A minimum of three separate animals were analyzed per condition, with each digit sourced from a separate animal.

#### Quantification of hyaluronic acid area after digit amputation

Using FIJI, brightness and contrast for all images were adjusted in batch to reduce background, with the same settings applied to each sample. The HABP fluorescence channel was segmented using default automatic thresholding settings. The specificity and sensitivity of the segmentation step was evaluated for each sample, and the total segmented area was calculated per field of view (FOV). Two sections per digit were analyzed, and these values were averaged and considered a single biological replicate. A minimum of three separate animals were analyzed per condition, with each digit sourced from a separate animal.

#### Morphometric quantification of digit length and area

Fixed digits were imaged on a Leica stereomicroscope (Leica, Wetzlar, Germany), and images were analyzed in FIJI, where the region of the nail was manually outlined as an ROI using the Polygon Selection tool, and the nail length was traced using the Straight Line tool. For the 4-MU experiment, each mouse was considered a biological replicate, and for experiments involving direct injections of substances, each digit was considered a biological replicate.

#### Quantification of the number of Ki67^+^ cells, SP7^+^ cells, and collagen area per FOV after 4- MU treatment versus control

For Ki67 quantification, IHC images were taken of 200 µm Z-stacks. ROIs of identical size were selected, specifically focusing upon the blastema in the control digits and the connective tissue overlying the P3 bone in the 4-MU digits (excluding the marrow). The Ki67 fluorescence channel was used to determine the total number of Ki67^+^ nuclei in the FOV. A minimum of 3 separate animals were analyzed per condition, with each digit sourced from a separate animal and considered as a single biological replicate. For SP7 quantification, ROIs included both the edge of the P3 bone and connective tissue overlying it. The SP7 fluorescence channel was used to determine the total number of SP7^+^ nuclei in the FOV. For collagen area, the ROI centered upon the soft connective tissue overlying bone. The SHG channel was automatically segmented, and area was calculated. For these samples, two sections per digit were analyzed, and these values were averaged and considered as a single biological replicate. A minimum of 3 separate animals were analyzed per condition, with each digit sourced from a separate animal and considered as a single biological replicate.

#### Quantification of *in situ* and *in vitro* pSMAD1/5/8 fluorescence intensity

For quantification of pSMAD1/5/8 fluorescence intensity of 4-MU versus control samples, all images were converted to 8-bit on FIJI. ROIs of identical size were created, focusing upon the connective tissue surrounding the P3 bone. The nuclei of the samples were automatically selected using the Hoechst channel, and all ROIs were saved. The same ROIs were then applied to the pSMAD1/5/8 channel, and average fluorescence intensity was calculated. A minimum of 3 separate animals were analyzed per condition, with each digit sourced from a separate animal and considered as a single biological replicate.

For quantification of pSMAD1/5/8 fluorescence intensity of cells cultured on hydrogels of different stiffnesses, all images were converted to 8-bit on FIJI. Then, ROIs were automatically created using the Hoechst channel to identify nuclei. Saved ROIs were applied to the pSMAD1/5/8 channel to calculate the fluorescence intensity. A minimum of three independent experiments were performed, and values for at least 10 separate cells were averaged as a single biological replicate.

#### Quantification of membrane CD44 signal intensity on hydrogels

Using FIJI, all images were converted to 8-bit. Automatic thresholding was performed on the CD44 channel to segment membrane CD44, and average signal intensity was calculated. A minimum of three independent experiments were performed, and intensity values for at least 10 separate cells were averaged as a single biological replicate.

#### Quantification of pericellular HA and HAPLN1 on hydrogels

Using FIJI, all images were converted to 8-bit. Thresholding was performed on the F-actin channel to segment the entire cell body, and the total area of the cell body was calculated. The ROI of the cell body was used to segment HA and HAPLN1 on the cell surface, and the area covered by HA and HAPLN1 were calculated. Pericellular HA and HAPLN1 was quantified as the division of HA or HAPLN1 area by cell body area. A minimum of three independent experiments were performed, and values for at least 10 separate cells were averaged as a single biological replicate.

#### Quantification of pericellular HA in mCherry Control versus Hapln1^OE^ cells

The HABP channel was used to automatically segment HA, and the percentage area containing HA was quantified per FOV. A minimum of three independent experiments were performed, and each hydrogel was imaged at 5 different locations, which were averaged as a single biological replicate.

#### Statistical analysis

Statistical analysis was performed using GraphPad PRISM (GraphPad Software, La Jolla, CA). The Shapiro-Wilk test was used to assess normality, and outliers were identified by ROUT tests (for multiple outliers, Q = 1%). A two-tailed student’s *t*-test was used for pairwise comparisons, and a two-way or three-way ANOVA followed by Tukey test was used for multiple comparisons. P < 0.05 was considered statistically significant. Data were presented as mean ± SEM.

---

**Figure S1.**
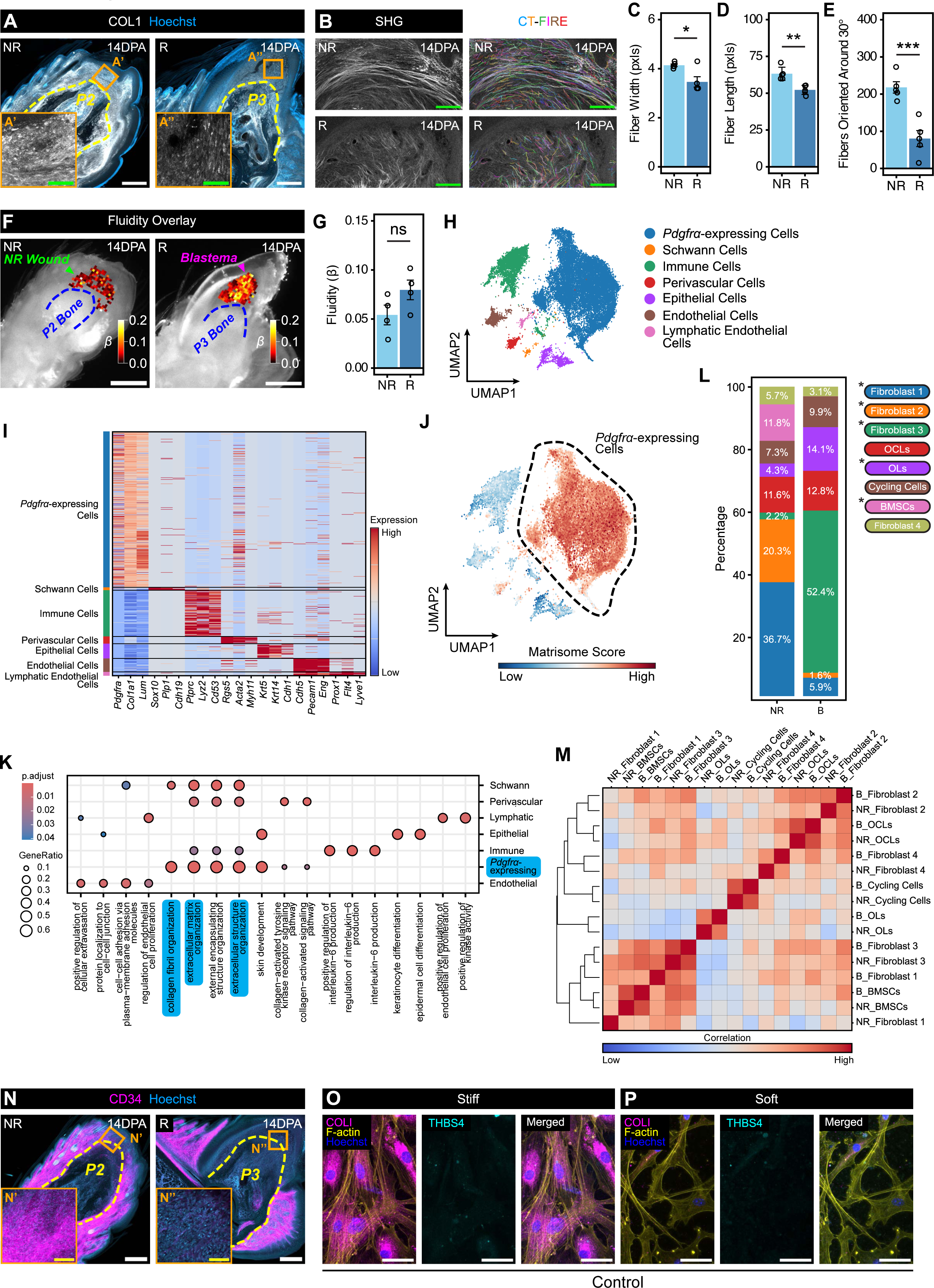
Digit wound healing responses diverge in their cellular and ECM composition, related to Figure 1. (A) Immunofluorescence images of 14 days post-amputation (DPA) non-regenerating (NR) and regenerating (R) digits stained for COLI (white) and Hoechst (blue). The yellow dashed line indicates the border of the second (P2) or third (P3) phalanx bone. Scale bars = 250 µm. Insets show the orange boxed regions of fibrosing (A’) and blastema (A”) tissues at higher magnification. Scale bars = 50 µm. (B-E) Second harmonic generation (SHG) images of 14DPA NR and R digits. Scale bars = 100 µm. Collagen fibers were segmented using curvelet transform fiber extraction (CT-FIRE), and fiber width (C), length (D), and alignment (E) were quantified. n = 5 mice per condition. (F and G) Atomic force microscopy fluidity maps of the 14DPA NR wound (green arrowhead) and blastema (magenta arrowhead). The blue dashed line indicates the border of the P2 or P3 bone. Scale bars = 500 µm. Average fluidities were quantified in (G), which were determined by applying the Power Law Rheology model. n = 4 mice per condition. (H) UMAP of the identities of seven major cell types in the wounded digit (combined non-regenerative and blastema datasets). They include *Pdgfrα*-expressing (blue), Schwann (orange), immune (green), perivascular (red), epithelial (purple), endothelial (brown), and lymphatic endothelial (pink) cells. (I) Heat map of top differentially expressed genes for each major cell population depicted in (H). (J) UMAP plot scoring each cell by their expression of matrisome genes, where dark red indicates high expression and dark blue low expression. The matrisome score was calculated as the average expression of the matrisome gene list subtracted by the average expression of a reference gene set. The *Pdgfrα*-expressing cluster (black dashed circle) scored the highest. (K) Gene ontology terms upregulated for each major cell population, determined by gene set enrichment analysis using clusterProfiler. The size of the circle corresponds to the proportion of input genes associated with a gene ontology term, and the color indicates the adjusted p-value of the enrichment score. (L) Cell proportions of *Pdgfrα*-expressing sub-types in the NR digit and blastema. Values embedded in the plot represent average proportions among all *Pdgfrα*-expressing cells within a wound healing condition. Statistical testing was performed using the *propeller* package, which employs moderated *t*-tests in an empirical Bayes framework. The asterisk (*p < 0.05) denotes significant differences in proportion between wound healing conditions. OCLs = osteochondral- lineage cells, OLs = osteo-lineage cells, and BMSCs = bone marrow stromal cells. (M) Correlation matrix comparing all fibroblast sub-types separated by NR and blastema conditions. Positive correlation is shown as red or negative correlation as blue. NR = non-regenerating, B = blastema, BMSCs = bone marrow stromal cells, OLs = osteo-lineage cells, OCLs = osteochondral-lineage cells. (N) Immunofluorescence images of 14DPA NR and R digits stained for CD31 (white), CD34 (magenta), and Hoechst (blue). The yellow dashed line indicates the border of the P2 or P3 bone. Scale bars = 250 µm. Insets show the boxed regions of fibrosing (N’) and blastema (N”) tissues at higher magnification. Scale bars = 50 µm. (O and P) Immunofluorescence of fibroblasts cultured on stiff (50 kPa; O) or soft (0.7 kPa; P) hydrogels without ascorbic acid and stained for COLI (magenta), F-actin (yellow), THBS4 (cyan), and Hoechst (blue). Scale bars = 50 µm. Data are shown as mean ± SEM. Statistical significance was determined by two-tailed unpaired student’s *t*-test. *p < 0.05, **p<0.01, ***p<0.001, ns = not significant.

**Figure S2.**
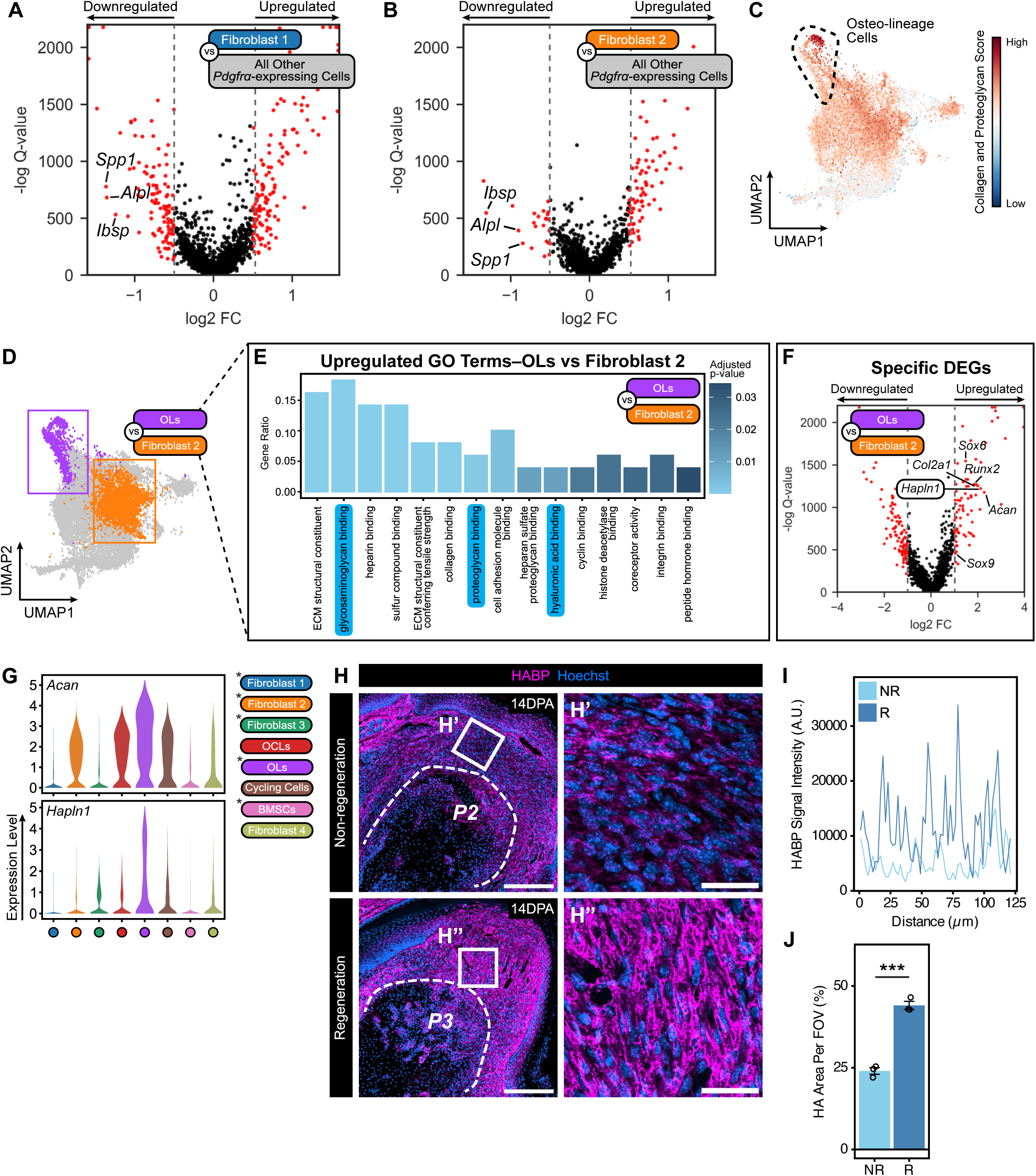
Osteo-lineage cells and hyaluronic acid binding characterize the blastema, related to Figure 2. (A and B) Volcano plots of differentially expressed genes in Fibroblast 1 (A) or Fibroblast 2 (B) cells compared to all other *Pdgfrα*-expressing cells, determined by the two-part generalized linear model MAST. Genes related to osteogenesis are annotated. Thresholds were set at -0.5 > log2FC > 0.5 and Q-value < 0.05. (C) UMAP plot scoring each cell by their expression of collagen and proteoglycan genes, where dark red indicates high expression and dark blue low expression. The osteo-lineage cells (black dashed circle) display the highest expression of these genes. (D) UMAP diagram showing the comparison of the osteo-lineage cluster (OLs, purple) against Fibroblast 2 cluster (orange). Boxed regions are highlighting the clusters of interest. (E) Gene ontology terms upregulated in osteo-lineage cells (OLs) compared to Fibroblast 2 cells, determined by gene set enrichment analysis using clusterProfiler. The Gene Ratio indicates the proportion of input genes associated with a gene ontology term, and the color indicates the adjusted p-value of the enrichment score. (F) Volcano plot of differentially upregulated genes in osteo-lineage cells (OLs) compared to Fibroblast 2 cells, determined by the two-part generalized linear model MAST. Genes related to hyaluronic acid and chondrogenesis are annotated. Thresholds were set at -1 > log2FC > 1 and Q-value < 0.05. (G) Violin plots of *Hapln1* and *Acan* expression by all *Pdgfrα*-expressing cell sub-types. OCLs = osteochondral-lineage cells, OLs = osteo-lineage cells, and BMSCs = bone marrow stromal cells. (H-J) Immunofluorescence images of 14 days post-amputation (DPA) non-regenerating (NR) or regenerating (R) digits stained for HABP (magenta) and Hoechst (blue). The white dashed line indicates the border of the second (P2) or third (P3) phalanx bone. Scale bars = 200 µm. The right-hand panels show higher magnification views of the white boxed regions of fibrosing (H’) and blastema (H”) tissues. Scale bars = 35 µm. Pericellular HA described by HABP signal intensity shown in arbitrary units (A.U.) across distance between non-regenerating (NR, light blue) and regenerating (R, dark blue) digits in (I). Quantification of the percentage of hyaluronic acid (HA) area per field of view (FOV) between 14DPA NR and R digits in (J). n = 3 mice per condition. Data are shown as mean ± SEM. Statistical significance was determined by two-tailed unpaired student’s *t*-test. *p < 0.05, **p<0.01, ***p<0.001, ns = not significant.

**Figure S3.**
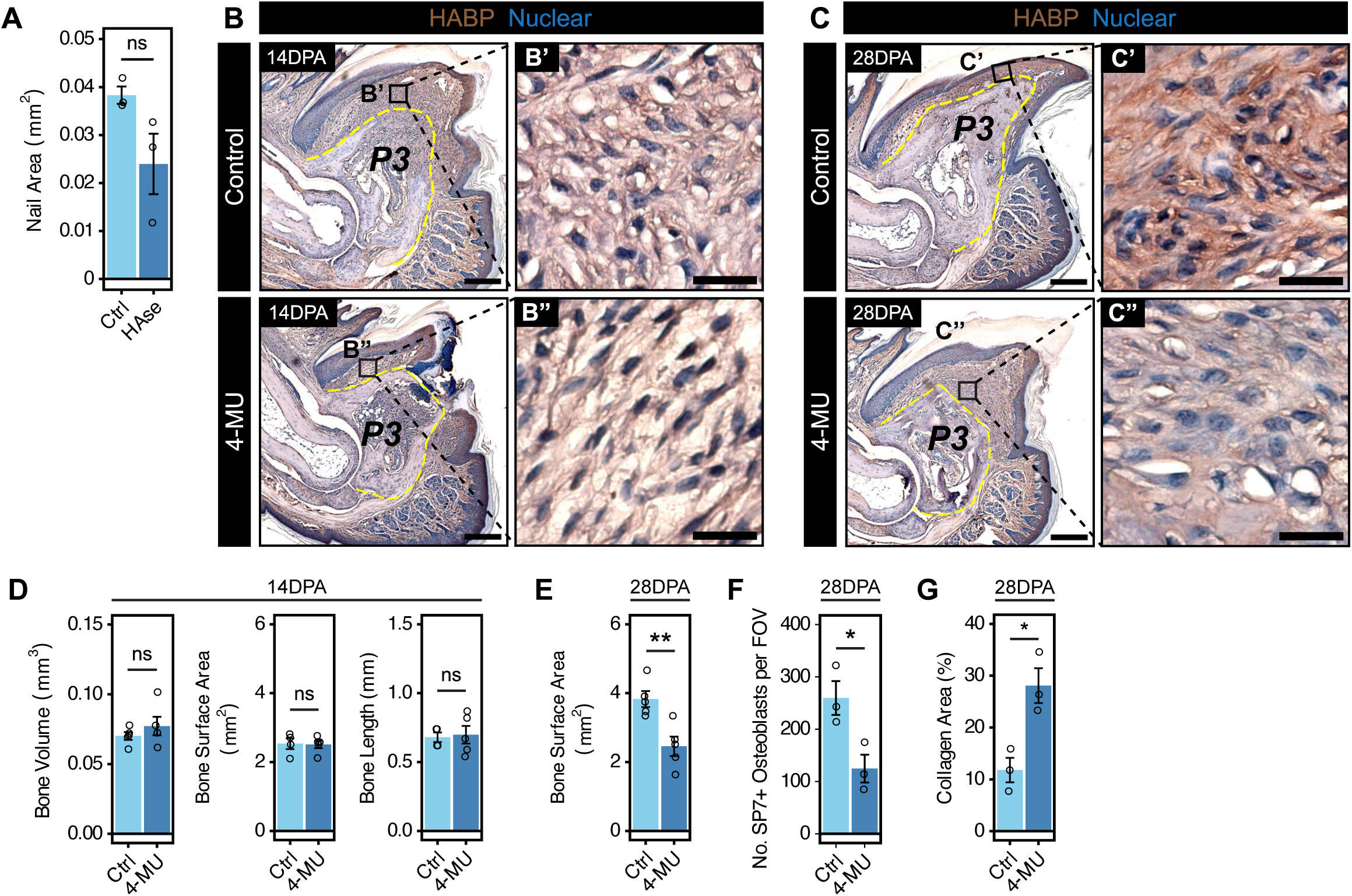
Hyaluronic acid depletion impairs digit tip regeneration, related to Figure 3. (A) Quantification of nail area of 14 days post-amputation (DPA) control and hyaluronidase (HAse) digits, related to Figure 3B. n = 3 mice per condition. (B and C) Brightfield chromogenic images of control (top panel) and 4-MU (bottom panel) digits at 14DPA (B) and 28DPA (C) stained for HABP and nuclear counterstained with hematoxylin. The yellow dashed lines indicate the border of the third phalanx (P3) bone. Scale bars = 250 µm. The right-hand panels show regions of the blastema (B’, C’) and 4-MU wound (B”, C”) at higher magnification. Scale bars = 25 µm. (D) Quantification of P3 bone volume (left), surface area (middle), and length (right) of 14DPA control and 4-MU digits, related to Figure 3J. n = 5 mice per condition. (E) Quantification of P3 bone surface area of 28DPA control and 4-MU digits, related to Figure 3K. n = 5 mice per condition. (F) Quantification of the number of SP7^+^ osteoblasts per field of view (FOV) in 28DPA control and 4-MU digits, related to Figure 3P. n = 3 mice per condition. (G) Quantification of the percentage collagen area in 28DPA control and 4-MU digits, related to Figure 3P. n = 3 mice per condition. Data are shown as mean ± SEM. Statistical significance was determined by two-tailed unpaired student’s *t*-test. *p < 0.05, **p<0.01, ***p<0.001, ns = not significant.

**Figure S4.**
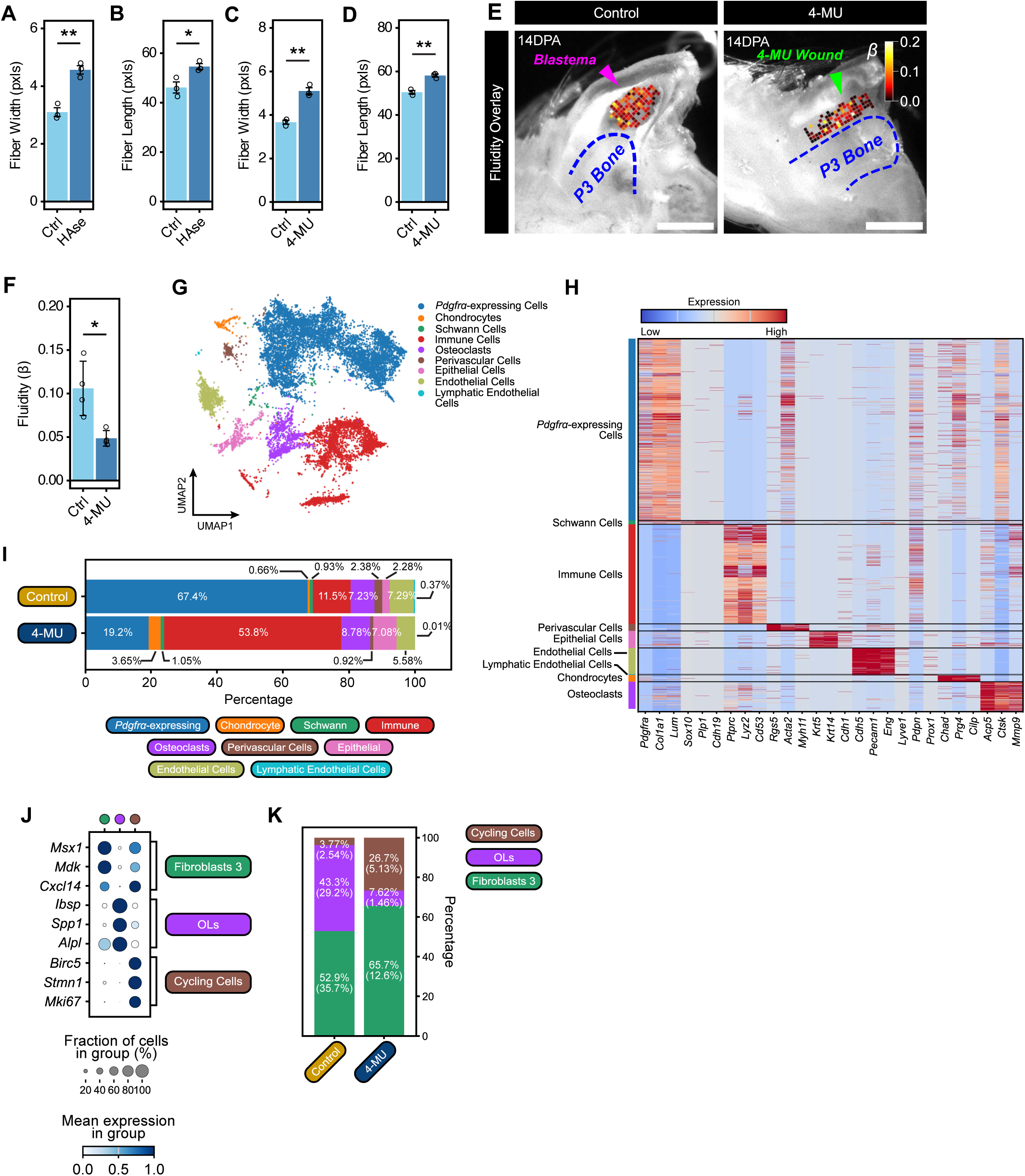
Perturbation of HA alters the amputated digit’s cellular and ECM composition, related to Figure 4. (A and B) Quantification of collagen fiber width (A) and length (B) in control and hyaluronidase (HAse) digits, related to Figure 4B. n = 3 mice per condition. (C and D) Quantification of collagen fiber width (C) and length (D) in control and 4-MU digits, related to Figure 4F. n = 3 mice per condition. (E and F) Atomic force microscopy fluidity maps of the 14 days post-amputation (DPA) control blastema (magenta arrowhead) and 4-MU wound (green arrowhead). The dashed blue line indicates the border of the third phalanx (P3) bone. Scale bars = 500 µm. Average fluidities were quantified in (G), which were determined by applying the Power Law Rheology model. n = 4 mice per condition. (G) UMAP of the identities of nine major cell types in the control and 4-MU digits at 14DPA. They include *Pdgfrα*-expressing (blue), chondrocytes (orange), Schwann (green), immune (red), osteoclast (purple), perivascular (brown), epithelial (pink), endothelial (yellow), and lymphatic endothelial (turquoise) cells. (H) Heat map of top differentially expressed genes for each major cell population that was identified in (G). Dark red indicates high gene expression and blue low expression. (I) Cell proportions of all cell types in control and 4-MU digits at 14DPA. Values embedded in the plots represent average proportions. (J) Dot plot showing top differentially expressed genes for each *Pdgfrα*-expressing cell sub-type. The size of the circle represents the fraction of cells expressing the gene, and average gene expression is shown according to the key, in which dark blue and white are high and low expression, respectively. OLs = osteo-lineage cells. (K) Cell proportions of *Pdgfrα*-expressing cell sub-types in control and 4-MU digits at 14DPA. Values embedded in the plots represent average proportions among *Pdgfrα*-expressing cells, and parenthetical values represent average proportions among all cells. OLs = osteo-lineage cells. Data are shown as mean ± SEM. Statistical significance was determined by two-tailed unpaired student’s *t*-test or two-way ANOVA with Tukey’s multiple comparisons test. *p < 0.05, **p<0.01, ***p<0.001, ns = not significant.

**Figure S5.**
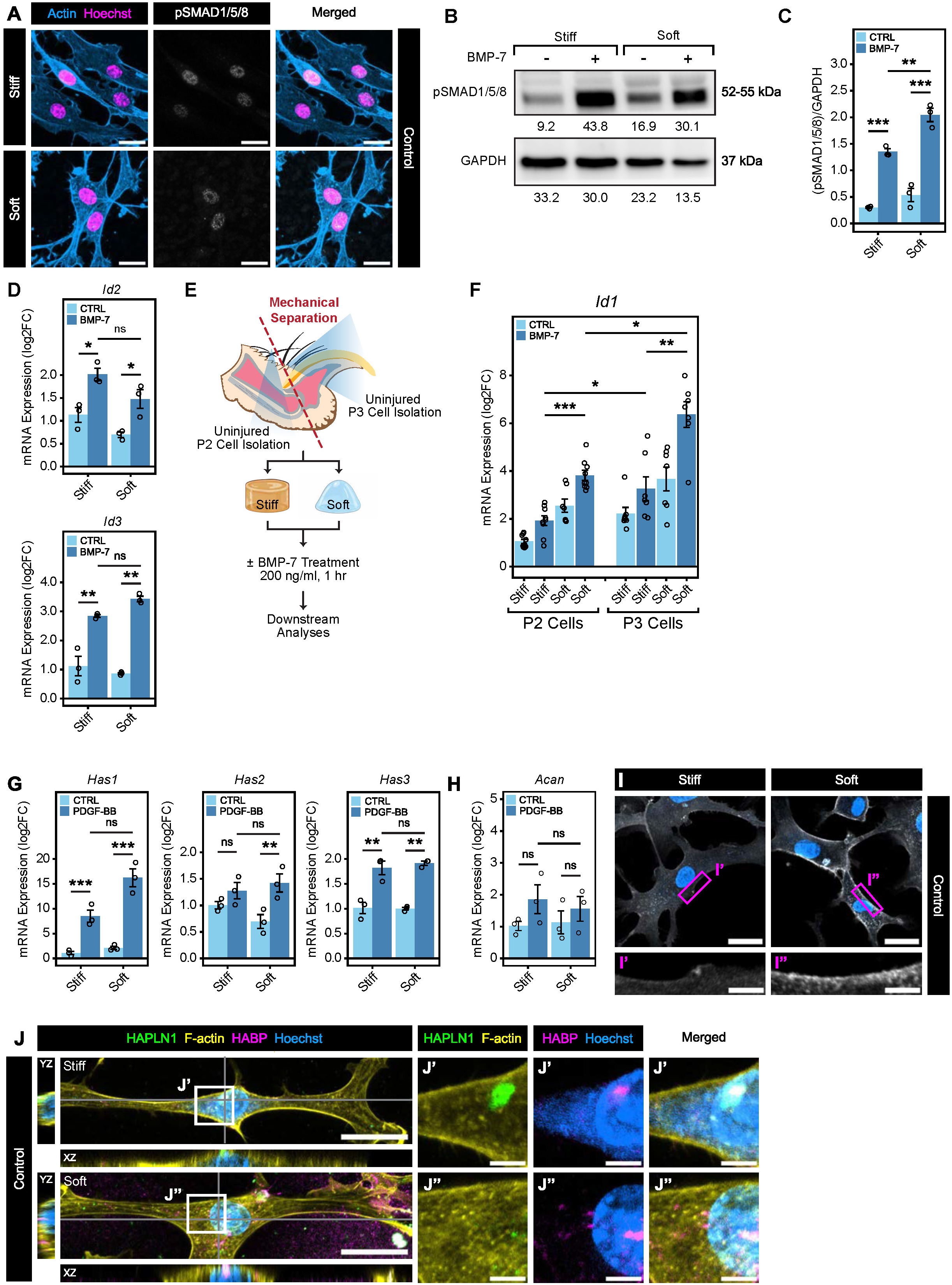
Soft substrates enhance BMP signaling, related to Figure 5. (A) Immunofluorescence images of fibroblasts cultured on stiff (50 kPa) or soft (0.7 kPa) hydrogels without BMP-7 and stained for pSMAD1/5/8 (white), F-actin (blue), and Hoechst (magenta). Scale bars = 20 µm. (B and C) Immunoblotting of pSMAD1/5/8 and GAPDH as the loading control in fibroblasts cultured on stiff (50 kPa) or soft (0.7 kPa) hydrogels, with or without BMP-7. Values below the blots represent the integrated band intensities. The integrated band intensities of pSMAD1/5/8 were normalized to GAPDH and represented in (C). n = 3 independent experiments. (D) qPCR analyses of *Id2* and *Id3* gene expression in fibroblasts cultured on stiff (50 kPa) or soft (0.7 kPa) hydrogels, with or without BMP-7. n = 3 independent experiments. (E) Schematic of the experimental design testing the influence of substrate stiffness on BMP signaling in second (P2) and third (P3) phalanx cells. (F) qPCR analysis of *Id1* gene expression in P2 or P3 cells cultured on stiff (50 kPa) or soft (0.7 kPa) hydrogels, with or without BMP-7. n = 7-9 independent experiments. (G and H) qPCR analyses of *Has1*, *Has2*, *Has3* (G) and *Acan* (H) gene expression in fibroblasts cultured on stiff (50 kPa) or soft (0.7 kPa) hydrogels, with or without PDGF-BB. n = 3 independent experiments. (I) Immunofluorescence images of fibroblasts cultured on stiff (50 kPa) or soft (0.7 KPa) hydrogels without PDGF-BB and stained for CD44 (white) and Hoechst (blue). Scale bars = 25 µm. The bottom panels show regions of the cell membrane (magenta boxes) at higher magnification. Scale bars = 10 µm. (I) (J) Immunofluorescence images of PDGF-BB control fibroblasts cultured on stiff (50 kPa) or soft (0.7 kPa) hydrogels and stained for HABP (magenta), HAPLN1 (green), F-actin (yellow), and Hoechst (blue). XZ and YZ images are orthogonal views at the plane of the gray solid lines. Scale bars = 25 µm. The right-hand panels show the white boxed regions (J’ and J”) at higher magnification in split images. Scale bars = 5 µm. Data are shown as mean ± SEM. Statistical significance was determined by two-way ANOVA with Tukey’s multiple comparisons test. qPCR data were normalized to Stiff-Control conditions and shown as log2FC, with statistical analyses performed on -11CT values. *p < 0.05, **p<0.01, ***p<0.001, ns = not significant.

**Figure S6.**
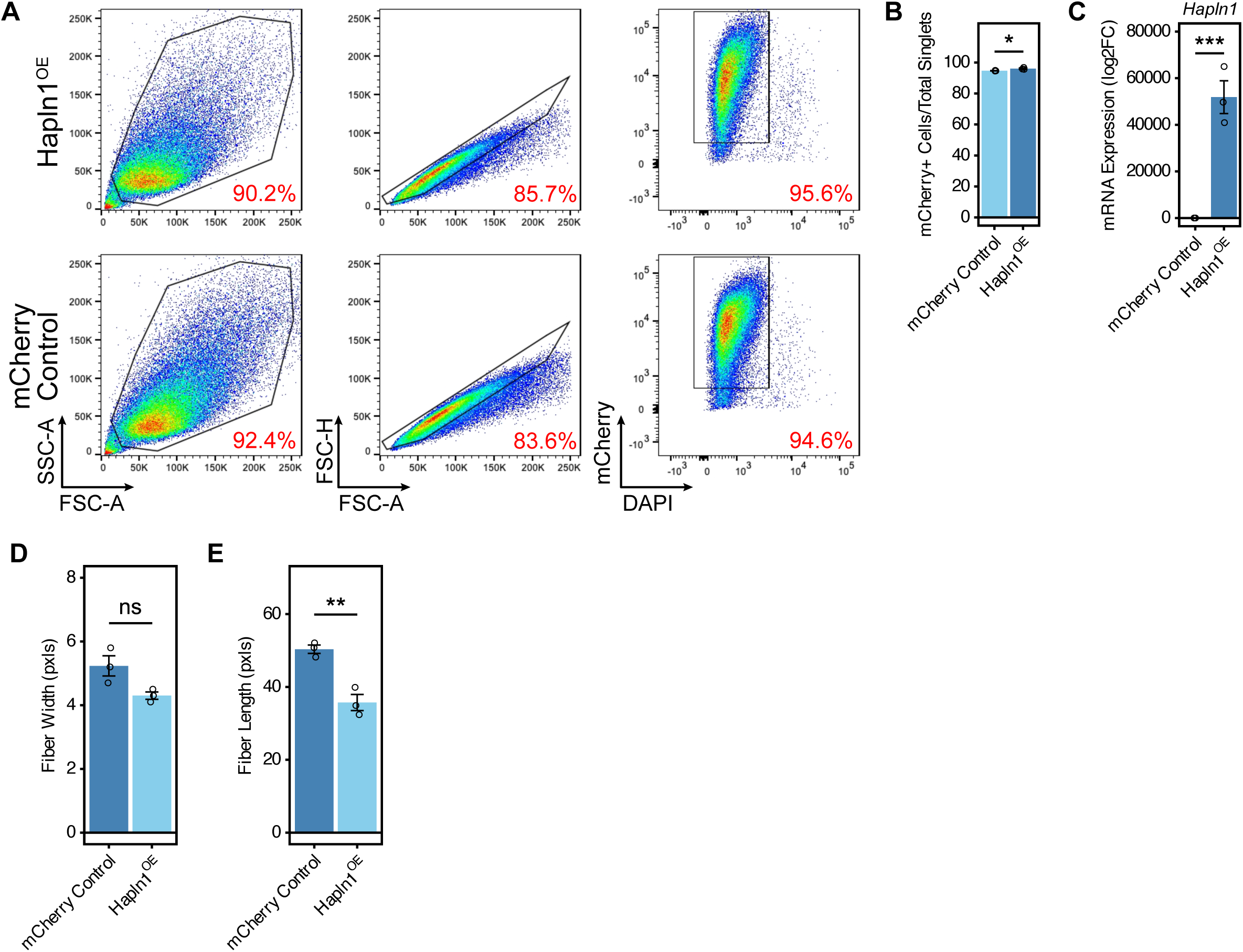
*Hapln1* overexpression disrupts collagen fibrillogenesis, related to Figure 6. (A and B) Flow cytometric analysis of transduced mCherry Control and Hapln1 overexpression (Hapln1^OE^) fibroblasts. The stepwise gating strategy involves exclusion of cell debris (FSC-A and SSC-A), followed by doublets (FSC-A and FSC-H), and finally dead, mCherry^-^ cells. mCherry Control and Hapln1^OE^ infection efficiencies were quantified in (B). n = 3 independent experiments. (I) (C) qPCR analysis of *Hapln1* gene expression in mCherry Control and Hapln1^OE^ fibroblasts. n = 3 independent experiments. (D and E) Quantification of collagen fiber width (H) and length (I) in mCherry control and Hapln1^OE^ fibroblasts using CT-FIRE, related to Figure 6H. n = 3 independent experiments. Data are shown as mean ± SEM. Statistical significance was determined by two-tailed unpaired student’s *t*-test. qPCR data were normalized to mCherry Control fibroblasts and shown as log2FC, with statistical analyses performed on -11CT values. *p < 0.05, **p<0.01, ***p<0.001, ns = not significant.

**Figure S7.**
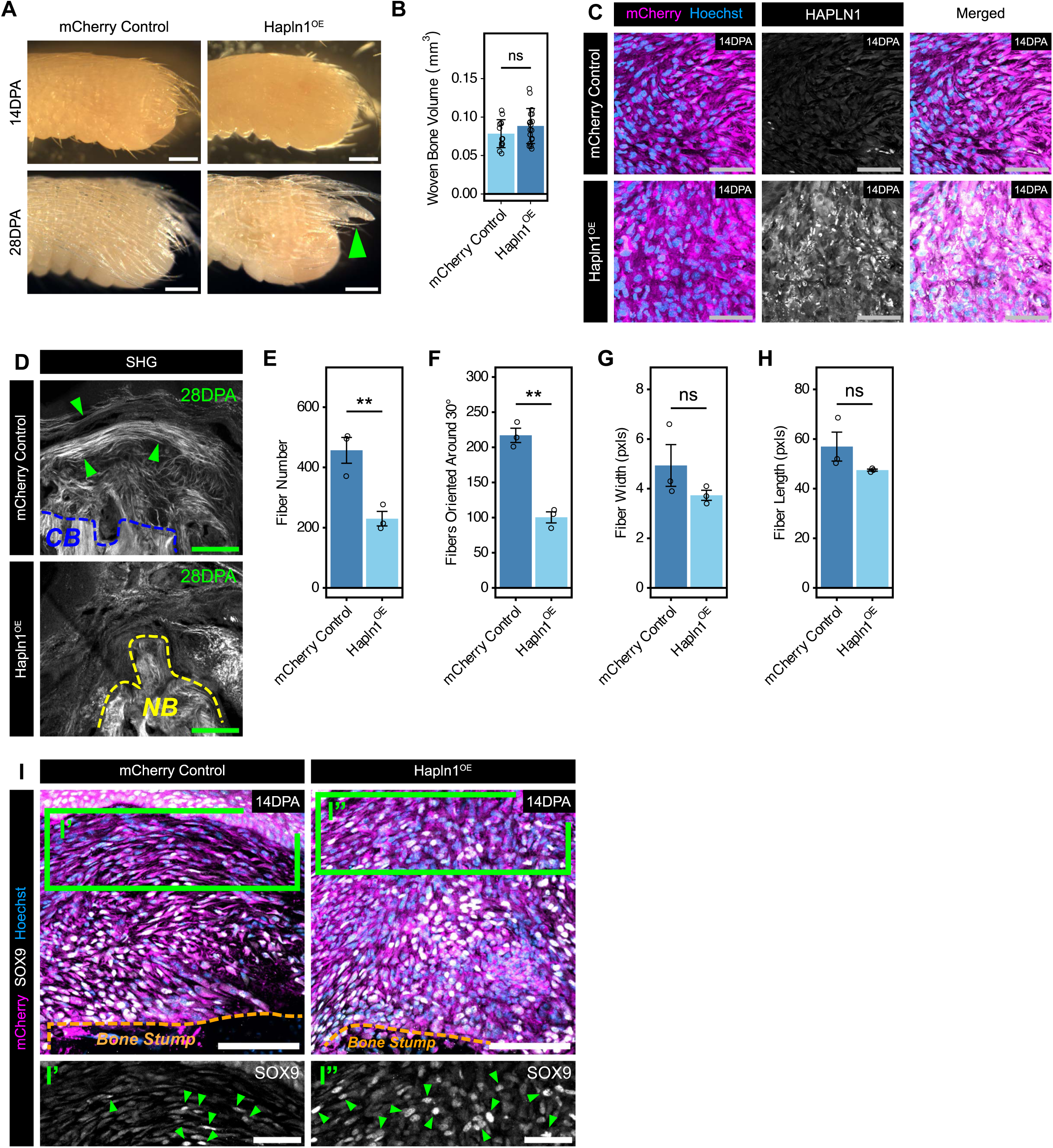
HAPLN1 promotes osteogenic differentiation and collagen remodeling, related to Figure 7. (A and B) Representative images of 14DPA (top panels) and 28DPA (bottom panels) digits after mCherry Control or *Hapln1* overexpressing (Hapln1^OE^) fibroblast transplantation. The green arrowhead indicates a sprouting nail. Scale bars = 150 µm. (B) Quantification of new bone volume after mCherry Control or Hapln1^OE^ fibroblast transplantation in 28DPA digits, related to Figure 7B. n = 18 digits. (C) Immunofluorescence images of 14DPA digits after mCherry Control or Hapln1^OE^ fibroblast transplantation and stained for mCherry (magenta), HAPLN1 (white), and Hoechst (blue). Scale bars = 50 µm. (D-H) Second harmonic generation (SHG) microscopy images of 28DPA digits after mCherry Control or Hapln1^OE^ fibroblast transplantation, highlighting collagen I fibers. The blue dashed line indicates the border of the pre-existing cortical bone (CB), and the yellow dashed line indicates the border of new bone (NB) formation. The green arrowheads indicate regions of collagen fibers. Scale bars = 125 µm. Fiber number (E), orientation (F), width (G), and length (H) were quantified using CT-FIRE. n = 3 mice per condition. (I) Immunofluorescence images of 14DPA digits after mCherry Control or Hapln1^OE^ fibroblast transplantation. The orange dashed line indicates the border of the second phalanx (P2) bone stump. Scale bars = 100 µm. Green boxed regions (I’ and I”) are shown at higher magnification in the bottom panels. Green arrowheads point at SOX9^+^ nuclei. Scale bars = 50 µm. Data are shown as mean ± SEM. Statistical significance was determined by two-tailed unpaired student’s *t*-test. *p < 0.05, **p<0.01, ***p<0.001, ns = not significant.

