## Supplemental Table 1 for "Hyaluronic Acid and Emergent Tissue Mechanics Orchestrate Digit Tip Regeneration"

| **Datasets** | **ncounts min filter** | **ncounts max filter** | **ngenes min filter** | **ngenes max filter** | **Mitochondrial percentage max filter** | **Scrublet filter score** | **Ambient RNA removal procedure** |
| --- | --- | --- | --- | --- | --- | --- | --- |
| Regen 14DPA MS 1 | 0 | 50000 | 500 | 7000 | 10% | 0.25 | N/A |
| Regen 14DPA MS 1 | 0 | 50000 | 600 | 7000 | 10% | 0.25 | N/A |
| Regen 14DPA Jessica | 0 | 70000 | 200 | 7000 | 10% | 0.25 | N/A |
| NonRegen14DPA 1 | 0 | 60000 | 500 | 4500 | 10% | 0.25 | N/A |
| NonRegen14DPA 2 | 0 | 75000 | 2200 | 8500 | 10% | 0.25 | SoupX |
| NonRegen14DPA 3 | 0 | 40000 | 2000 | 6000 | 10% | 0.2 | SoupX |
| 4MU Control 1 | 1000 | 20000 | 400 | 5000 | 10% | 0.2 | Cellbender |
| 4MU Control 2 | 4000 | 20000 | 600 | 6000 | 10% | 0.2 | Cellbender |
| 4MU Treat 1 | 1000 | 25000 | 400 | 6000 | 10% | 0.2 | Cellbender |
| 4MU Treat 2 | 1000 | 50000 | 1500 | 6000 | 10% | 0.2 | Cellbender |
