## Supplemental Table 2 for "Hyaluronic Acid and Emergent Tissue Mechanics Orchestrate Digit Tip Regeneration"

| **Table 2. Recipe for fabricating hydrogels of different stiffnesses** | | | | | | | |
| --- | --- | --- | --- | --- | --- | --- | --- |
| ***E* (kPa)** | **40% Acrylamide (µl)** | **2% Bis-acrylamide (µl)** | **2 M AHA (µl)** | **TEMED (µl)** | **10% APS (µl)** | **MiliQ H_2_O (µl)** | **Total Volume (µl)** |
| 0.7 | 35 | 27.2 | 20 | 2.5 | 5 | 410.3 | 492.5 |
| 50 | 94 | 70.5 | 20 | 2.5 | 5 | 308 | 492.5 |
